# Enhanced behavioural and neural sensitivity to punishments in chronic pain and fatigue

**DOI:** 10.1101/2024.04.04.588151

**Authors:** Flavia Mancini, Pranav Mahajan, Anna á V. Guttesen, Jakub Onysk, Ingrid Scholtes, Nicholas Shenker, Michael Lee, Ben Seymour

## Abstract

Chronic pain and fatigue in musculoskeletal disease contribute significantly to disability, and recent studies suggest an association with reduced motivation and excessive fear avoidance. In this behavioural neuroimaging study in chronic inflammatory arthritis participants and healthy controls, we aimed to identify the specific behavioral and neural changes associated pain and fatigue during reward and loss decision-making. Computational modeling of behaviour identified a parametric signature, characterized most notably by increased punishment sensitivity. This signature is distinct from patterns previously reported in psychiatric conditions and it aligns with predictions of mechanistic models of chronic pain such as the fear avoidance model. Neural activity associated with the punishment prediction error was enhanced in the right posterior insular cortex, putamen, pallidum, and dorsolateral prefrontal cortex. Functional network connectivity analysis showed that insula centrality correlated with subjective reports of fatigue and pain. Overall, the findings show that pain and fatigue in chronic pain relate to objective behavioural changes, and can be mapped to a specific pattern of activity in brain circuits of motivation and decision-making.

## 1 INTRODUCTION

Chronic pain and fatigue are the cardinal symptoms of musculoskeletal disease and lead to a level of disability that has significant socioeconomic cost (Woolf and Pfleger, 2003). Although multiple factors may contribute to the risk of developing chronic pain, a key hypothesis is that reduced central motivational drive to engage in physical activity leads to physical deconditioning that in turn exacerbates pain, a corollary of the fact that physiotherapy is clearly beneficial in most subacute and chronic musculoskeletal conditions (Meulders, 2019; Vlaeyen et al., 2016). This ‘fear avoidance’ hypothesis links with other neurobiological and cognitive factors known to contribute to risk for pain chronification, embedded within the broader biopsychosocial model that captures the complexity of chronic pain in everyday clinical contexts (Gatchel et al., 2007). The mechanism by which pain reduces central motivation therefore represents a critical link in the pathway to chronic pain and fatigue development (Seymour et al., 2023), but its nature and neurobiology are not known.

Injuries are common across animal species, for example after accidents and within/cross species contests, and lead to recognisable protective and recuperative behaviours (Walters and Williams, 2019). These are presumed to be evolutionarily adaptive because the injured state makes animals more vulnerable to further harm, and so being extra careful whilst recovering benefits survival (Walters, 1994). Peripheral and spinal sensitization to pain provide one route to hyper-protective behaviour, and excessive pain hypersensitivity is a key focus of many current models of chronic pain. But recent ecological studies indicate a broader hyper-sensitivity to threat that goes beyond pain, suggesting a central, supramodal enhanced sensitivity to punishments as a mechanism of behavioural homeostasis after an injury, and a potential contributing factor to chronic pain and disability (Lister et al., 2020). Whilst enhanced punishment sensitivity would be adaptive for simple musculoskeletal injuries encountered in ecological contexts, it would ultimately be maladaptive in the context of clinical musculoskeletal conditions where inactivity is detrimental (Seymour et al., 2023).

Computational models of the neural circuits of learning and motivation allow us to dissect its core information processing steps, thus helping identify the underlying components of post-injury homeostasis such as modulation of punishment sensitivity (Seymour and Mancini, 2020). A similar approach has been applied to human behavioural experiments to identify, for example, reduced reward sensitivity in major depressive disorder, and aversive generalisation in anxiety disorder (Aylward et al., 2019; Huys et al., 2013; Dunsmoor and Paz, 2015). In this study, we aimed to identify i) whether there is a specific behavioural signature of pain and fatigue manifest during reward and loss decision making, and ii) if there is, does it relate to specific, localisable activity in the brain, in particular in circuits associated with motivation and learning.

Reward and loss decision-making captures a fundamental process that underlies much of our everyday behaviour - doing things that benefit us, and avoiding things that don’t. We studied this using a 4-armed bandit task (figure 1; (Daw et al., 2006; Seymour et al., 2016; Aylward et al., 2019)). Computational modelling of behavioural data can then be used to interrogate brain activity to find corresponding neural processing steps, and their relation to pain and associated fatigue. In the human brain, the mesolimbic and insula regions are thought to play a key role in pain-related motivation and avoidance learning (Fazeli and Büchel, 2018; Horing and Büchel, 2022; Navratilova and Porreca, 2014; Geuter et al., 2017), alongside a broader role in behavioural homeostasis (e.g. in illness and sickness states (Uddin et al., 2017)). We reasoned that the insula in particular might link the physiological identification of injury with responsiveness to punishments, which may ultimately contribute to the pain and fatigue phenotype.

**Figure 1.**
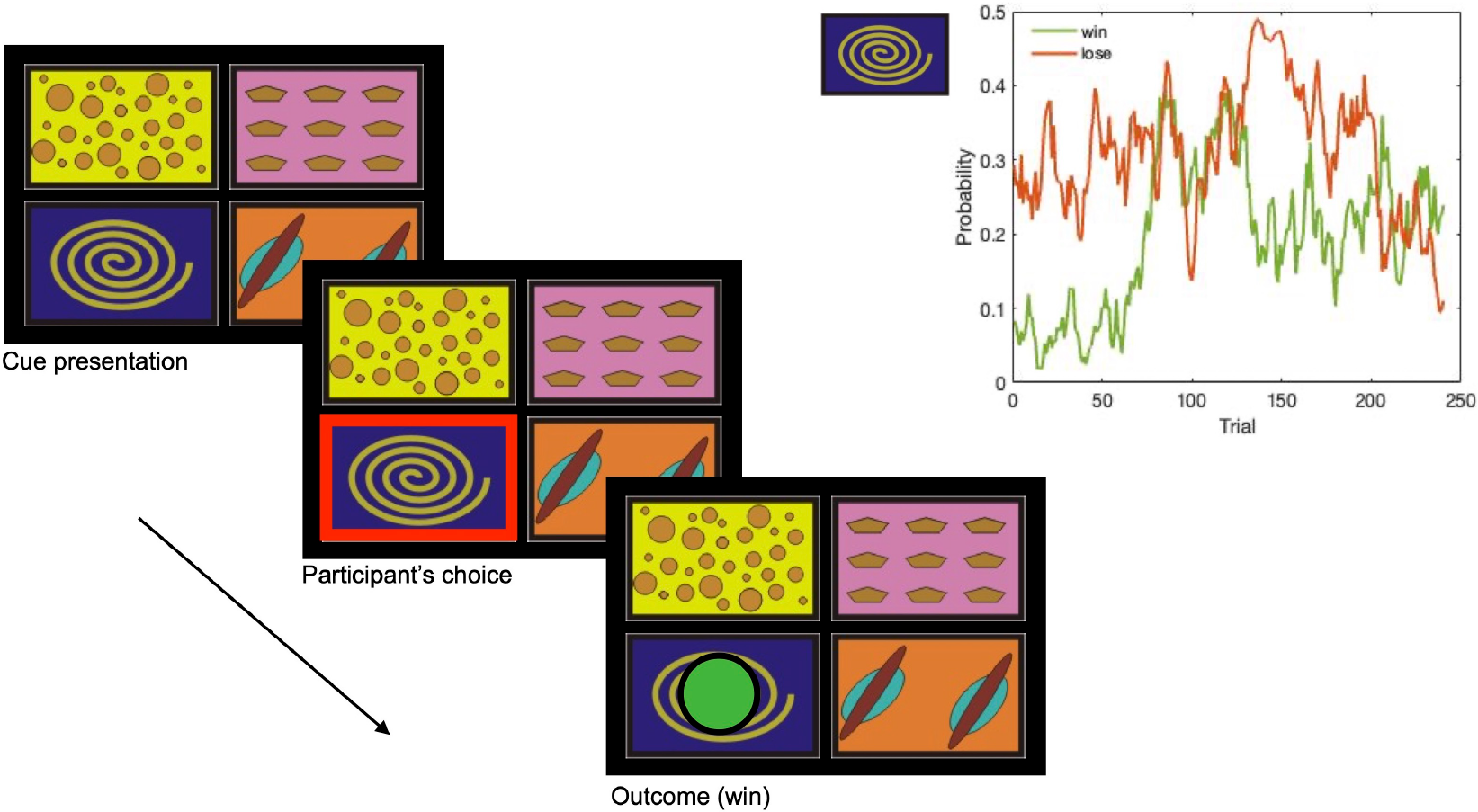
Task design. At the beginning of each trial, four cues were presented and participants chose one of them. Each cue was associated with an independent probability of winning and losing a token. After each choice, the outcome was displayed (green coin = win £1).

## 2 RESULTS

### 2.1 Choice behaviour

We recruited 60 volunteers to a behavioural neuroimaging experiment: 30 patients with painful inflammatory arthritis, and 30 age and sex matched controls (see Methods). In the 4-armed bandit task, participants were presented with four cues, and they needed to choose one of them on each trial. Each cue was associated with an independent, non-stationary probability of winning or losing a coin worth £1. The mean reward and punishment rate were independent of each other, and noisy - slowly changing across trials as a Gaussian random walk. After selecting one of the cues, their choice yielded one of four possible outcomes: one green token (win £1), one red token (lose £1), one green and one red token (win £1 but lose £1, so effectively £0 when added at the end of the game), or no outcome. Thus, participants need to constantly and independently learn about rewards and losses, allowing us to dissociate differences in reward and loss learning.

Basic performance measures (net gains, reaction times, frequency of choice switches) were comparable between patients and controls (see Supplementary Behavioural Results). This was expected: as the task is effectively a dynamic learning environment for probabilistic outcomes, different levels of performance do not yield big differences in outcomes, which protects against the potentially confounding effects of accruing different winnings. In contrast, any differences between groups is embedded in the trial-by-trial dynamics as a result of exploration and learning.

To capture these learning effects we used a hierarchical Bayesian inference approach to fit multiple variants of Reinforcement Learning (RL) models to the choice data, first to all pooled participants and then separately for the two groups (see Methods). These are biological informed models, arising from basic animal learning theory, that are known to capture behaviour and brain activity during this general class of task (Seymour and Mancini, 2020). The hierarchical Bayesian approach allows for individual differences while aggregating information across participants of the same group and make meaningful group comparisons (Ahn et al., 2017). The choice data of the pooled participants and of the patient group (Table 1) were best fitted by models 1 and 3. These two models included two sensitivity parameters for rewards and punishments, which were used to scale the prediction error, a lapse and a decay parameter capturing forgetting in choice behaviour. In absolute terms, model 1 fitted best the data, but it was not significantly different than model 3. The only difference between models 1 and 3 designs lied in their learning rates: model 1 had two learning rates (*alpha* and *beta*) that were added for updating reward values (reward LR = *α* + *β*) and subtracted for updating punishment values (punishment LR = *α*-*β*), whereas model 3 had separate, fixed learning rates for rewards and punishments. In the control group, model 3 showed the best absolute fit, but it was not significantly different than the other models we tested.

**Table 1.** Model comparison for the 4-armed bandit task. Five models were fitted to all participants, as well as the arthritis and control groups separately: (1) differential learning rates (LR), lapse and decay parameters; (2) differential LR with lapse parameter, no decay; (3) separate LRs for rewards and punishments, lapse and decay parameters; (4) separate LRs for rewards and punishments, lapses but no decay; (5) separate LRs for rewards and punishments, without lapse or decay parameters. We report the expected log predictive density (ELPD) difference between the best performing model (lowest LOOIC (leave-one-out cross-validation information criterion)) and each model are displayed, alongside the standard error (SE) of the difference and the sigma effect (the ratio between the ELPD and SE difference, which is a proxy for significance).

| Group | Model | ELPD difference | SE difference | Sigma effect | LOOIC |
| --- | --- | --- | --- | --- | --- |
| All | 1 | 0 | 0 |  | 24386.3539 |
|  | 3 | -53.422223 | 37.8139003 | -1.4127668 | 24493.1983 |
|  | 2 | -276.00621 | 90.2032862 | -3.0598243 | 24938.3663 |
|  | 4 | -280.33323 | 91.9252409 | -3.0495785 | 24947.0203 |
|  | 5 | -336.24668 | 97.0027097 | -3.4663638 | 25058.8472 |
| Arthritis | 1 | 0 | 0 |  | 12754.9737 |
|  | 3 | -46.954621 | 37.2017901 | -1.2621603 | 12848.8829 |
|  | 4 | -221.65393 | 80.4995507 | -2.7534803 | 13198.2815 |
|  | 2 | -226.78714 | 77.0709632 | -2.9425757 | 13208.5479 |
|  | 5 | -264.26755 | 83.3235728 | -3.171582 | 13283.5088 |
| Controls | 3 | 0 | 0 |  | 11667.014 |
|  | 1 | -2.6956325 | 9.89283891 | -0.2724832 | 11672.4052 |
|  | 4 | -46.800276 | 48.2061096 | -0.970837 | 11760.6145 |
|  | 2 | -48.534824 | 49.9225293 | -0.9722028 | 11764.0836 |
|  | 5 | -56.156688 | 49.6040047 | -1.1320999 | 11779.3273 |

Our main interest was to evaluate group differences of model parameters. Hence, we compared the group estimates of the parameters of model 1 between the arthritis patient group and the control group, to investigate any core underlying differences in choice behaviour.

The group-level estimates of each free parameter of model 1 are reported in Table 2, and the individual participant’s estimates are shown in Figure 2. In keeping with our core hypothesis, participants with chronic arthritis were more sensitive to punishments when compared to controls, i.e. showed enhanced punishment sensitivity (P). In addition, we also found that they had faster memory decay and greater lapses in the choice behaviour. Finally, the differential component of the learning rate (*β*) was also higher in patients.

**Table 2.** Group-level parameter estimates for the arthritis and control groups (mean and SD) in model 3 (separately fitted for each group) and HDI group differences (lower and upper 95% HDI). Significant differences do not include 0 and are highlighted with *.

| Parameter | Arthritis | Controls | Lower 95% HDI | Upper 95% HDI |
| --- | --- | --- | --- | --- |
| reward sensitivity | 12.750 (5.911) | 9.417 (5.031) | -1.004 | 7.695 |
| punishment sensitivity | 9.953 (2.577) | 5.778 (3.715) | 1.388* | 8.016* |
| $\alpha$ | 0.585 (0.300) | 0.569 (0.178) | -0.133 | 0.183 |
| $\beta$ | 0.106 (0.100) | 0.016 (0.018) | 0.0130* | 0.101* |
| reward learning rate ($\alpha+\beta$) | 0.679 (0.069) | 0.590 (0.043) | -0.067 | 0.25 |
| punishment learning rate ($\alpha-\beta$) | 0.547 (0.074) | 0.570 (0.047) | -0.199 | 0.142 |
| lapse $\xi$ | 0.105 (0.043) | 0.026 (0.003) | 0.036* | 0.105* |
| decay | 0.603 (0.311) | 0.469 (0.261) | 0.008* | 0.403* |

**Figure 2.**
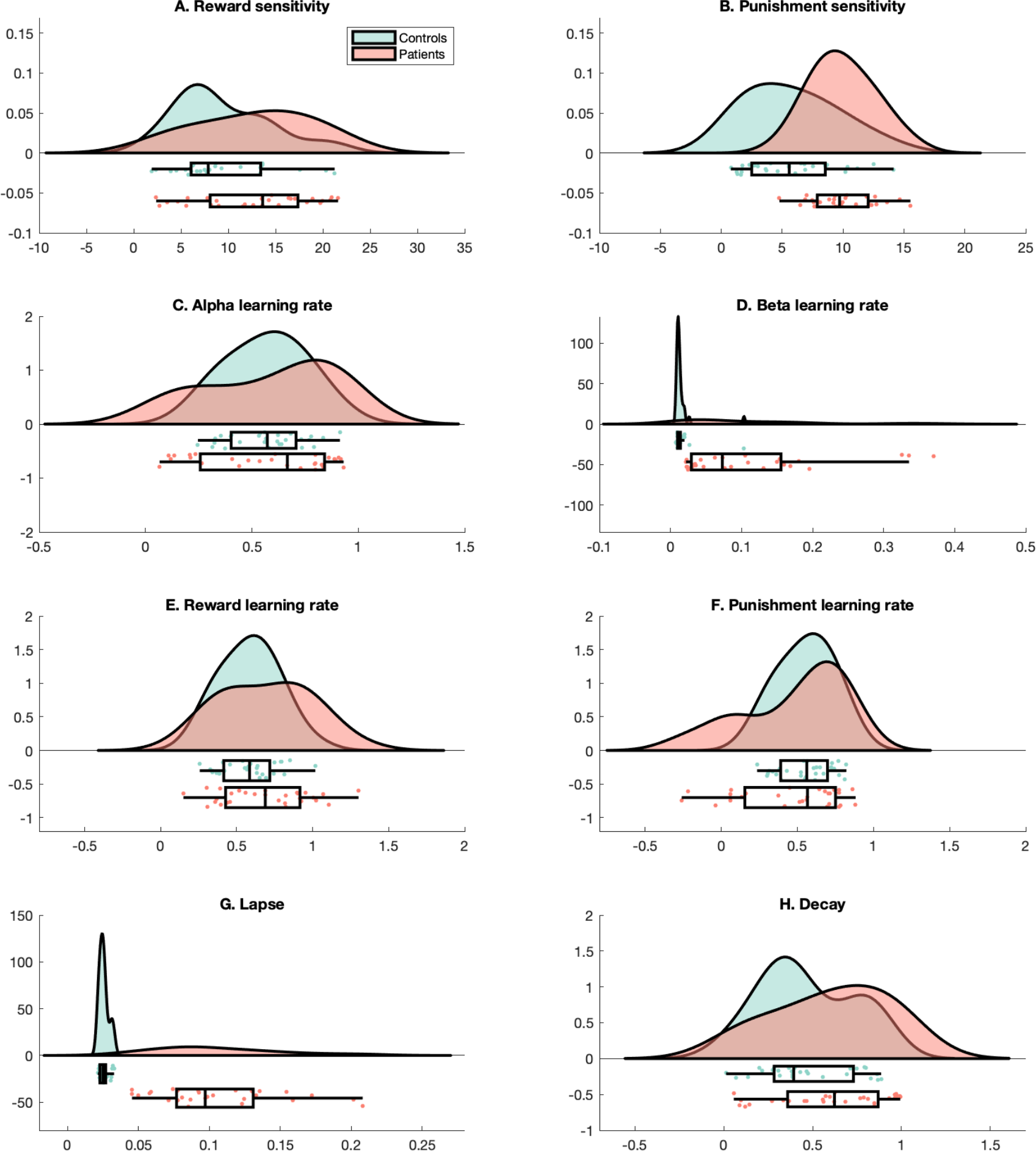
Participant’s estimates of each parameter. For each parameter (a.u.), we show the raincloud plots which include the (half) probability distribution function, scatterplots and boxplots of the data of arthritis patients (red) and controls (green).

### 2.2 Neural correlates of punishment PE, PPE

Participants performed the bandit task whilst undergoing functional brain MRI. We aimed to identify brain changes associated with reward and loss decision making. Since punishment sensitivity scales the magnitude of the value of the punishment as used for learning avoidance actions, this is reflected in brain responses correlated with the punishment prediction error (PPE), with the individually estimated punishment sensitivity as a covariate. Thus, we evaluated whether the neural activity associated with the PPE, and modulated by punishment sensitivity, differed across groups. The arthritis group showed increased activity in the right posterior insular cortex, putamen, pallidum, and dorsolateral prefrontal cortex (figure 3; table 3. There was no significant cluster of decreased activity, at the cluster threshold used (z>2.6, p<0.05). For completeness, common activity across the two groups is reported in Supplementary Figure S3 and Supplementary Table S6.

**Table 3.** Activation clusters of increased activity in arthritis participants vs controls, associated with PPE and modulated by the punishment sensitivity parameter. Coordinates in MNI space.

| Cluster Id | Voxels | P | Z peak | X | Y | Z |
| --- | --- | --- | --- | --- | --- | --- |
| 2a | 808 | 0.0015 | 4.28 | 36 | -6 | -6 |
| 2b |  |  | 3.78 | 28 | -4 | -4 |
| 2c |  |  | 3.72 | 16 | 4 | 2 |
| 2d |  |  | 3.51 | 52 | 10 | 6 |
| 2e |  |  | 3.47 | 24 | 12 | -8 |
| 2f |  |  | 3.23 | 50 | -8 | 10 |
| 1a | 468 | 0.0282 | 4.1 | 28 | 46 | 32 |
| 1b |  |  | 3.51 | 14 | 18 | 34 |
| 1c |  |  | 3.49 | 16 | 22 | 30 |
| 1d |  |  | 3.39 | 16 | 32 | 32 |

**Figure 3.**
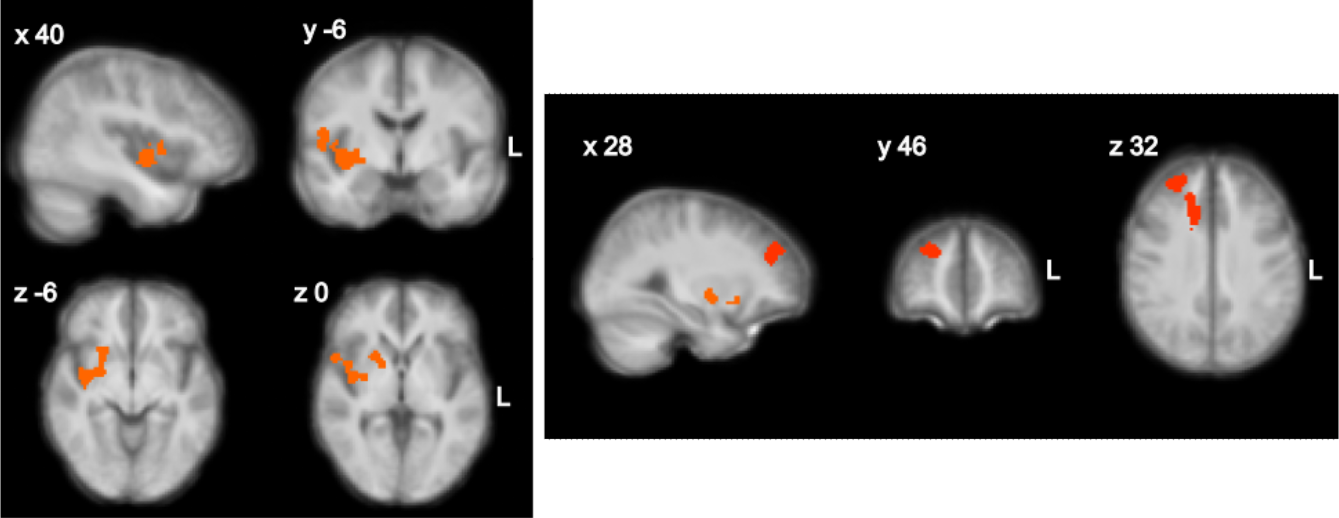
Increased neural processing of the Punishment Prediction Error in the arthritis group vs. controls, modulated by punishment sensitivity. Increased BOLD responses associated with PPE in the right posterior insula, putamen, pallidum and the dorsolateral prefrontal cortex. The colorscale represents z scores ranging from 0 to 5, thresholded at z>2.6, p<0.05. The statistical contrast was overlaid over a structural brain image, obtained by averaging the high-resolution structural images of the study participants.

**Figure 4.**
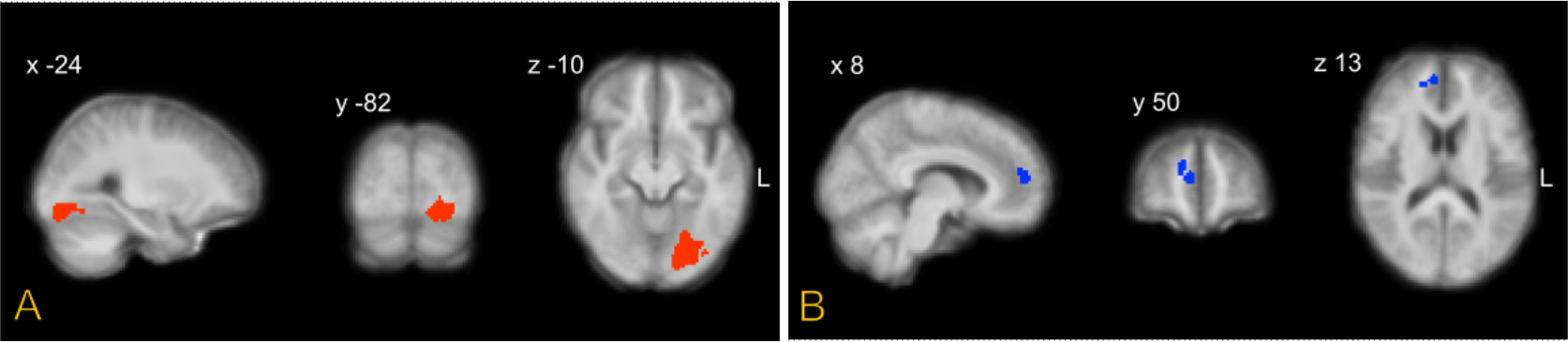
Increased (red, A) and decreased (blue, B) activity associated with the reward prediction error in the arthritis group vs. controls. Colorscale represents z scores ranging from 0 to 5, thresholded at z>2.6, p<0.05. The statistical contrast was overlaid over a structural brain image, obtained by averaging the high-resolution structural images of the study participants.

### 2.3 Neural correlates of reward PE, RPE

Rewards prediction error (RPE) activity modulated by reward sensitivity was decreased in arthritis participants vs. controls in the a rostral region of the paracingulate gyrus and neighbouring white matter, and increased in the left occipital fusiform gyrus, area V4 (table 4). There was no significant cluster of increased activity, at the cluster threshold used (z>2.6, p<0.05). Common activity across groups is reported in Supplementary Figure S4 and Table S7.

**Table 4.** Activation clusters of increased and decreased activity in arthritis patients vs controls, associated with RPE and modulated by the reward sensitivity parameter. Coordinates in MNI space.

| Cluster Id | Voxels | P | Z peak | X | Y | Z |
| --- | --- | --- | --- | --- | --- | --- |
| Arthritis > Controls |  |  |  |  |  |  |
| 1a | 662 | 1.31e-06 | 3.76 | -24 | -82 | -10 |
| 1b |  |  | 3.75 | -16 | -82 | -12 |
| 1c |  |  | 3.21 | -14 | -88 | -6 |
| 1d |  |  | 3.11 | -28 | -74 | -8 |
| 1e |  |  | 2.89 | -20 | -64 | -8 |
| 1f |  |  | 2.81 | -34 | -78 | -12 |
| Arthritis < Controls |  |  |  |  |  |  |
| 1a | 218 | 1.65 | 3.59 | 16 | 44 | 22 |
| 1b |  |  | 2.72 | 8 | 50 | 14 |
| 1c |  |  | 2.48 | 6 | 52 | 20 |

### 2.4 Connectivity analyses

The correlation of punishment sensitivity with evoked BOLD responses in the insula is in keeping with the core hypothesis that the insula is a central part of a behavioural homeostatic network that mediates motivational changes in the pain state. To look at this further, we aimed to identify whether the insula connectivity during the task correlates with pain behaviour and task parameters. To do this, we calculated the network centrality (here, degree centrality), which captures the importance or influence of the insula as a network hub during the task. We parcellated the brain into 180 symmetrical ROIs using a functional (Glasser) brain atlas, and constructed a binarised connectivity network using thresholded pairwise Pearson correlations (see Methods). This analysis ignores the task events, and simply considers overall connectivity during the task as a whole. We found that degree centrality in posterior insula subregions correlated with subjective reports of daily average pain and fatigue scores (see figure 5A). There was evidence for a correlation between Posterior Insular Area 2 and measures of pain (average daily pain rating from weekly diaries: Pearson’s r= 0.566, LogBF10= 3.413; BPI severity score: Pearson’s r= 0.544, LogBF10= 2.958), as well as fatigue (average daily fatigue rating from weekly diaries: Pearson’s r= 0.517, LogBF10= 2.468). These correlations were less apparent at rest, i.e. when performing the same analysis using the resting state fMRI data rather than task fMRI data (figure 5B). This has also been observed in previous work (Zhao et al., 2023).

**Figure 5.**
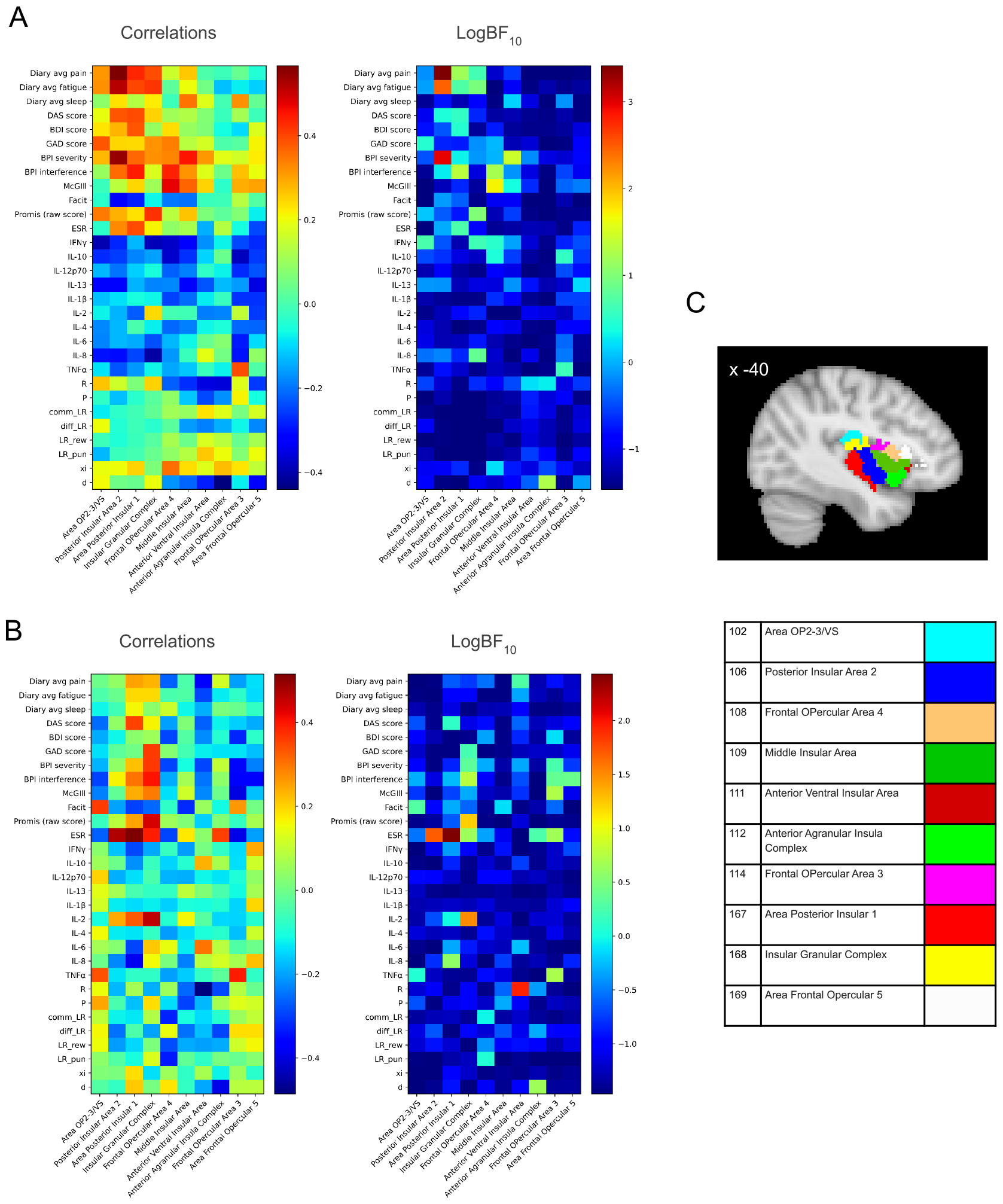
(A) Bayesian Pearson correlation of degree centrality of 10 insular nodes with clinical outcomes and model parameters for the patient group in task fMRI. (B) Bayesian Pearson correlation of degree centrality of 10 insular nodes with clinical outcomes and model parameters for the patient group in resting-state fMRI Significant correlations: Area Post Insular 1 with ESR (Pearson’s r= 0.515, LogBF10= 2.43) (C) A colour-coded map of insular nodes derived from the Glasser atlas (number denotes the parcel number in the Glasser atlas). Results for other network centralities are provided in Figs.S5, S6, S7, S8.

The analysis of connectivity during instrumental learning provides evidence that posterior insula activity not only correlates with pain related changes in objective task behaviour, but that it sits within a network of regions whose activity correlates with clinical subjective ratings of fatigue and pain.

## 3 DISCUSSION

The results reveal a specific computational signature of chronic pain and fatigue, reflected in the pattern of changes of the underlying parameters that govern motivated behaviour. At the heart of this signature is an increase in punishment sensitivity, which demonstrates a generalised form of hyper-protective behaviour that enhances learning of aversive outcomes and may lead to excessive avoidance. This allows the organism to enhance defence beyond transient pain, which can mediated by central and peripheral hypersensitivity, to other aversive outcomes.

The signature also includes other components of behaviour, including the learning rate differential between rewards and punishments, alongside increased forgetting and choice lapses. This reveals a specific pattern that is distinct from that seen in other clinical conditions, using the same or similar task. For instance, depression is associated with the opposite tendency, a reduced sensitivity to rewards (Huys et al., 2013)). Anxiety is associated with increased punishment learning rates (Aylward et al., 2019), opposite to the relative increase in reward learning rate seen here, suggesting more dynamic updating of rewards compared punishments. This is equivalent to a more persistent punishment memory, as it has been demonstrated in previous studies of fibromyalgia - a non-inflammatory pain condition with dominant pain and fatigue (Meulders et al., 2017). The increased sensitivity to punishments in inflammatory arthritis is also different from the reduced general outcome sensitivity observed in Parkinson’s disease as the result of deep brain stimulation (Seymour et al., 2016) and the reduced reward outcome value reported following trypotophan depletion (Seymour et al., 2012). An increase in the lapse and decay rates was also found in Parkison’s patients, as in our study. This adds randomness to the decisions and probably reflects a non-specific, disruptive effect of pain and fatigue on attention. Together, this illustrates how the chronic pain and fatigue state seen in inflammatory arthritis is clearly computationally distinct from depression, anxiety and Parkison’s disease.

The behavioural manifestation of computational components of learning and decision making maps well to the predictions of chronic pain models. Notably, increased punishment sensitivity is a central prediction of the the fear avoidance model (Meulders, 2019; Vlaeyen et al., 2016). Previous investigations have focused on punishment generalisation (Meulders et al., 2015), which is a distinct component of avoidance in which aversive values spread to perceptually similar pain predictors. Importantly, both increased punishment sensitivity and aversive generalisation drive the same tendency towards enhanced avoidance. This is a critical component of mechanistic models of chronic pain, which posit that the brain forms a hierarchical internal representations of the injury to drive protective and recuperative behaviours (Seymour and Mancini, 2020). Although greater avoidance reduces the chance of worsening damage through new actions, it inherently affects the opportunity to learn if an injury has resolved (information restriction), and will tend to perpetuate internal representations of injury states (Seymour et al., 2023). Future longitudinal studies would need to determine whether punishment sensitivity is a manifestation of the pain state or an outcome predictor.

At a cortical level, neural activity associated with punishment prediction error (PPE) was enhanced in the right posterior insular cortex, putamen, pallidum, and dorsolateral prefrontal cortex (dlPFC, figure 3) in the inflammatory arthritis group, according to individual differences in punishment sensitivity. The PPE reflects the change of punishment value at each timepoint that is used for learning, which is then scaled by the punishment sensitivity constant. In previous instrumental learning studies on healthy participants, PPE activity was reported in the anterior insula (Gueguen et al., 2021), putamen (Seymour et al., 2007), dlPFC (Gueguen et al., 2021). The role of the insula is in keeping with the hypothesis that it plays a fundamental role in pain (Horing and Büchel, 2022; Isnard et al., 2011; Moayedi, 2014; Segerdahl et al., 2015) and fatigue states (Stefanov et al., 2020; Wylie et al., 2020). This can also be related to a proposed role in active inference of interoceptive (Barrett and Simmons, 2015; Fermin et al., 2022), somatosensory (Allen et al., 2016) and nociceptive signals (Fazeli and Büchel, 2018; Geuter et al., 2017; Horing and Büchel, 2022).

The insula is well placed to interface afferent nociceptive information with efferent control through connectivity to mesolimbic and mesocortical pathways, other cortical sites involved in learning and motivation (such as the anterior cingulate and medial prefrontal cortex) and descending pain control systems. In our connectivity analysis, we computed the network centrality to capture the importance of different insula subregions as a network hub, and this was evidenced by the the correlation between posterior insula area 2 activity and subjective reports of daily fatigue and pain scores. We also note that this network metric was much more apparent during task performance than at rest. Although this finding needs to be cross-validated in new datasets, it might suggest that looking at brain connectivity in the context of motivated behaviour may be more sensitive than pure resting state brain activity, when used for biomarker generation. We should also note that we restricted our analysis to the insula, based on our a priori hypothesis, but it may well be that other regions (such as PFC and ACC) show symptom-correlated network activity.

Clinically, pain and fatigue are often highly correlated in inflammatory chronic pain conditions. However, the methods used in this study could be applied to painless, fatigue-dominant disorders, such as chronic fatigue syndrome and long COVID, to dissect the effect of fatigue from that of pain on instrumental learning. In inflammatory arthritis, the two are nearly impossible to disentangle. The approach here provides a behaviourally and computationally informed approach to disentangling these in future studies across other pain and fatigue cohorts.

## 4 METHODS

### 4.1 Participants

We initially engaged 15 local patients with chronic pain to help us design the study and iterated preliminary versions of the protocol using a systems engineering approach. The study was approved by the central NHS Health Research Authority ethics committee (REC: 17/SW/0113, IRAS: 216259). After the design was finalised, we screened 80 volunteers and recruited 30 patients with chronic rheumatoid and psoriatic arthritis and 30 healthy age-matched controls. Patients were recruited at Rheumatology, Cambridge University Hospitals. All participants gave informed written consent according to the Helsinki Guidelines. Three participants did not complete the full study and were therefore excluded, leaving us with 57 participants in total (28 controls and 29 patients; 45 females). Patients were 53.83 ± 8.09 years old (range 36-66 years; 27 females), and controls were 51.50 ± 10.77 years old (range 33-69 years; 18 females).

Participants were screened with a non-standardized demographic questionnaire and the following standardized questionnaires: Beck Depression Inventory II, Brief Pain Inventory, McGill Pain Assessment, Functional Assessment of Chronic Illness Therapy Fatigue Scale, Short form PROMIS Sleep Scale. Trained staff carried out arthritis assessments with standardized joint assessment disease activity score (DAS), rheumatoid classification (ACR/EULAR criteria) and psoriatic classification. Patients with disease activity (DAS >2.6) or subjectively high fatigue scores (>4 on scale 0 = ‘no fatigue’ to 10 = ‘worst fatigue imaginable’, FACIT <35) were included in the study. Controls had no clinically significant disease activity (DAS <2.6) or subjective fatigue (<4 on scale 0 = no fatigue to 10 = worst fatigue imaginable, FACIT >30). Participants with other explanations for fatigue (anaemia, unstable thyroid, heart conditions, diabetes, recent cancer treatment, recent surgery) and contra-indications for MRI safety (WBIC safety questionnaire) were considered not eligible to the study. The demographic and clinical details of the participants who completed the study are reported in Supplementary Table 1.

### 4.2 Procedure

Participants took part in an initial screening and training session, and in a further MRI session. The two sessions were separated by 5-73 days, depending on participant, site and scanner availability. Throughout both sessions and in between tasks/runs, participants were asked to rate their fatigue and pain on a scale of 0 to 10 (0 = no fatigue/pain, 10 = worst fatigue/pain imaginable).

### 4.3 4-armed bandit task

Participants underwent a 4-armed bandit decision-making task during MRI, following some practice. In each trial, participants could choose between 4 abstract boxes presented on the screen and try to win green tokens and simultaneously, but independently, try to avoid red tokens. The green and red signalled winning and loosing £1, respectively. The sum of positive minus negative tokens was exchanged for money at the end of the experiment. The probability of win/loss in each box changes gradually over trials as a gaussian random walk. The participants completed 240 trials over 4 runs.

### 4.4 Inflammatory markers

Blood samples were obtained for all participants in the second visit and the following inflammatory markers were evaluated: ESR, IFN*γ*, IL-10, IL-12p70, IL-13, IL-1 *β*, IL-2, IL-4, IL-6, IL-8, TNF *α*. General health was assessed in full blood count and liver function samples.

### 4.5 Actigraphy and daily ratings

Participants recorded daily ratings and wore an ActiGraph GT9X Link (Actigraph, LLC) to the wrist, 24 hours a day, over the course of 5-7 days. The wearable device contains a piezo-electric sensor that generates a voltage when the device undergoes a change in acceleration (3 axial). It was set to record 60s epochs. Participants removed the wristbands only for showering or swimming. During this week, participants rated their fatigue and pain (“how much fatigue/pain do you feel right now on a scale of 0 to 10?", whereby 0 = no fatigue/pain, 10 = worst fatigue/pain imaginable). The participants set their alarms in the morning (9:00/ 9:30) and evening (19:30/20:00) and noted their current rating down on a questionnaire that was previously provided by the experimenters. Participants also recorded their daily consumption of caffeine and alcohol on a questionnaire. They noted how many units were consumed and the details of the product (e.g. filter coffee, paracetamol, wine etc). Analysis and results of actigraphy is reported in the Supplementary Methods and Results document.

### 4.6 Modelling of choice behaviour

We simulated and fitted five Reinforcement Learning (RL) models based on non-probabilistic delta rules, whereby the learning rates are fixed and driven by discrepancies between the estimate of the expected value and observed values. In all models, we tracked the *Q* value of the four cues, as the sum of the *Q* values for rewards (*Q*^*r*^) and punishments (*Q*^*p*^). We calculated the reward and punishment prediction errors as the difference between the expected and observed value (Eq. 1, 2):

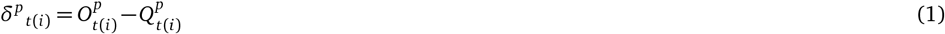

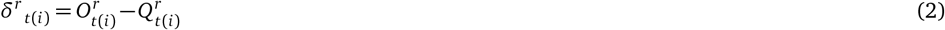

In all models, the prediction error was scaled by a free, sensitivity parameter (reward sensitivity *R* or punishment sensitivity *P*) (Eq. 3, 4), as in previous studies (Aylward et al., 2019):

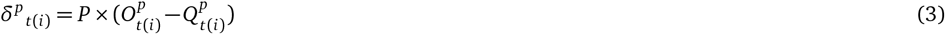

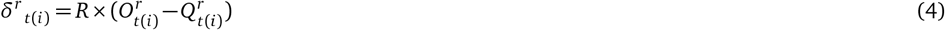

The models differed also according to the update rule, and the presence of lapses and forgetting in choice behaviour. In all models the learning rate was constant across trials. In models 1 and 2, the learning rate for rewards was calculated as the sum of two free parameters, *α* + *β*, and the learning rate for punishments was calculated as the difference of these free parameters, *α-β* :

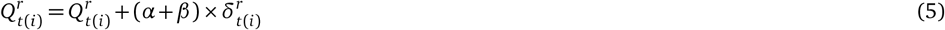

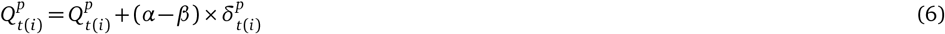

In models 3-5, the learning rates for rewards and punishments were estimated as two independent free parameters, i.e. *α*^*r*^ and *α*^*p*^:

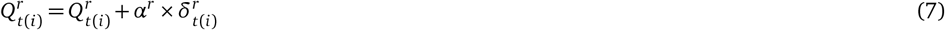

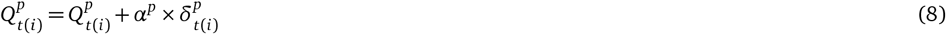

Models 1 and 3 additionally had a decay free parameter *d* to describe the rate of forgetting in the tracking of *Q*^*r*^ and *Q*^*p*^ (Eq. 9, 10), as in (Aylward et al., 2019):

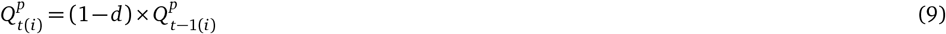

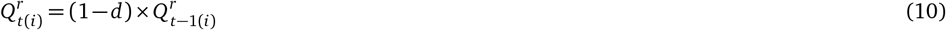

All models used a softmax policy to determine choice probability ℙ by combining the Q values of rewards and punishments:

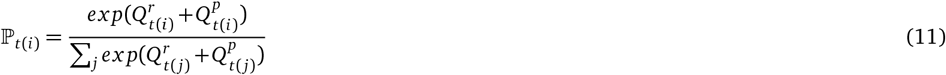

Models 1-4 softmax policy included a free parameter *ξ* that characterised lapses in choice behaviour, similarly to (Aylward et al., 2019):

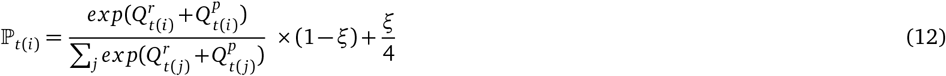

Table 5 summarises the free parameters of the different models we simulated and fitted to the data. We used hierarchical Bayesian methods (Hamiltonian Monte Carlo) to estimate model parameters, with the HBayesDM and Stan package for R (Stan Development Team, 2019; Ahn et al., 2017) in R. For each model, we fit 4 chains with 1000 burn in samples and 2000 samples to the following data: choice (1:4), win (0,1) and loss (0,-1) for each trial.

**Table 5.** Free parameters of the models that were fitted to the behavioural data.

| Model ID | Sensitivity | Reward LR | Punishment LR | Decay | Lapses |
| --- | --- | --- | --- | --- | --- |
| 1 | R, P | $\alpha + \beta$ | $\alpha - \beta$ | d | $\xi$ |
| 2 | R, P | $\alpha + \beta$ | $\alpha - \beta$ | n/a | $\xi$ |
| 3 | R, P | $\alpha^r$ | $\alpha^p$ | d | $\xi$ |
| 4 | R, P | $\alpha^r$ | $\alpha^p$ | n/a | $\xi$ |
| 5 | R, P | $\alpha^r$ | $\alpha^p$ | n/a | n/a |

Parameters for all models were fit separately for each group, as well as for all participants pooled in a common group. The winning model was defined as the model with the lowest Leave-One-Out Information Criterion (LOOIC). We also estimated the difference in models’ expected predictive accuracy through the difference in expected log point-wise predictive density (ELPD) (Vehtari et al., 2017). By taking the ratio between the ELPD difference and the standard error (SE) of the difference, we obtained the sigma effect, which is a heuristic for significance of such model differences.

We compared parameter estimates from the winning model across the two groups using 95 % high density intervals (HDI). In particular, for each comparison, we calculated the difference in the hyper parameters between groups and reported the 95 % HDI of the difference. In the Bayesian scenario, a significant group difference is indicated by the interval not containing the value 0.

### 4.7 MRI data acquisition

First, we collected a T1-weighted MPRAGE structural scan (voxel size 1 mm isotropic) on a 3T Siemens Magnetom Prisma scanner (Siemens Healthcare), equipped with a 64-channel head coil, at the Wolfson Brain Imaging Centre, Cambridge. Then we collected a resting state scan of 240 volumes and 4 task fMRI sessions of 264 volumes using the same multiband echo planar imaging (EPI) sequence (TR = 2000 ms, TE = 30 ms, flip angle = 82 deg, slices per volume = 64, multiband acceleration factor 2, interleaved slice mode, Grappa 2, voxel size 2 mm isotropic, A>P phase-encoding; this included four dummy volumes, in addition to those pre-discarded by the scanner). In order to correct for inhomogeneities in the static magnetic field, we imaged 4 volumes using an EPI sequence identical to that used in task fMRI, inverted in the posterior-to-anterior phase encoding direction. Full sequence metadata are available XX.

### 4.8 MRI data analyses

#### 4.8.1 Preprocessing

The anatomical images were reoriented to standard orientation, automatically cropped and bias-field corrected using ‘fsl_anat’ (FMRIB’s Software Library, www.fmrib.ox.ac.uk/fsl). The brain was extracted with ANTs tool ‘antsBrainExtraction’ (https://github.com/ANTsX/ANTs). Functional images had the first four volumes removed, were motion corrected using FSL tool ‘mcflirt’, 3D time shifted using AFNI program ‘3dTshift’ and a Fourier method (https://afni.nimh.nih.gov/). Fieldmap correction was applied using HySCO, Hyperelastic Susceptibility Artefact Correction (Ruthotto et al., 2013). Non-brain data were removed using FSL tool BET. Spatial smoothing was applied using a Gaussian kernel of FWHM 8.0mm. The entire 4D dataset was grand-mean intensity normalised by a single multiplicative factor and highpass temporal filtering was applied (Gaussian-weighted least-squares straight line fitting, with sigma=50.0s) using FSL.

#### 4.8.2 Generalised Linear Modelling

First and second level GLM analyses were conducted using FEAT (FMRI Expert Analysis Tool) Version 6.00, part of FSL. All univariate imaging results were obtained from a single GLM model. We investigated neural correlates using model 1. All model predictors were generated with the group mean fitted parameters in order to minimise noise. First level regressors included onset times and values for the reward outcomes, punishment outcomes, reward prediction error, punishment prediction error (modelled at the time of the outcome) and regressors of no interest, such as the cue onset and keypress time. Sessions within subject were not concatenated, but modelled separately and then averaged using a mixed effect model (second-level analysis). Third level analyses were Two-Group Difference models, in which we compared the positive and negative means of reward or punishment prediction errors between the arthritis and control groups; as covariates, we included the sensitivity parameter and age, demeaned across all subjects. Finally, we applied a cluster threshold z*>*2.6, p*<*0.05 and we report only clusters that survived this multiple-comparison correction.

#### 4.8.3 Connectivity analyses

Filtered functional images, preprocessed as per section 4.8.1 were registered to standard format (MNI template) and then parcellated into 180 regions of interest using the Glasser functional atlas (Glasser et al., 2016). Each region of interest combines the symmetric region in the brain, i.e. right and left regions jointly form one parcel. The timeseries for each parcel were extracted as the mean of the timeseries’ corresponding to all voxels in that parcel. This process was performed for all four runs of task-fMRI and one run of rest-fMRI, and we performed elementwise summation of the four task-fMRI timeseries runs, making it comparable with the rest-fMRI.

Fully connected networks were constructed with edge strength between two nodes being the Pearson correlation between the timeseries corresponding to the nodes. We then thresholded the Pearson’s full correlation matrices to a produce binary adjacency matrix (consisting of 1’s and 0’s) for each participant, as done in our previous studies(Mano et al., 2018). Each of the correlation matrices was thresholded in an adaptive manner to produce an adjacency matrix with a 10% link density. This value was chosen based on previous network topology studies that have found such a link density to provide optimal discriminative ability (Mano et al., 2018; Achard et al., 2012; Itahashi et al., 2014; Mansour et al., 2016; Termenon et al., 2016). We then computed the following nodal centrality measures for parcels corresponding to insular sub-regions: degree centrality, eigen-vector centrality, betweenness centrality, closeness centrality and load centrality.

The Glasser’s atlas, being a functional atlas, has several parcels, some of which may overlap with the anatomical insular region. We found the Glasser parcels which correspond to the anatomical insula region by cross-checking the regions which intersect/overlap with the left and right insula regions in the AAL atlas, in FSLeyes. This resulted in 10 parcels of the Glasser atlas, reported in Figure 5C with their corresponding area descriptions from the Glasser atlas. Then, we performed Bayesian Pearson correlation of insular nodal centrality metrics with clinical scores and model parameters, only for the patient group. We did not include any healthy subjects because they have normal clinical scores, and no pain, and correlations would have risked being confounded by group differences.

We correlated the nodal centrality metrics of these insular subregions with several scores and model variables: Daily average pain, Daily average fatigue, Daily average sleep quality, DAS/DAPSA score, BDI score, GAD score, BPI severity, BPI interferance, McGill questionnaire score, Facit score, Promis (raw score), Immunological variables (ESR, IFNg, IL-10, IL-12p70, IL-13, IL-1b, IL-2, IL-4, IL-6, IL-8, TNFa) and model parameters.

## 5 DATA AND CODE AVAILABILITY

All code and data will be available open source, released upon acceptance of the paper.

## Supporting information

supplementary information and analyses

## 6 ACKOWLEDGEMENTS

The study was funded by a Versus Arthritis (21537) grant to B.S. and M.L., a MRC Career Development Award to FM (MR/T010614/1) and a UKRI Advanced Pain Discovery Platform grant to both F.M. and B.S. (MR/W027593/1). B.S. was also funded by Wellcome (214251/Z/18/Z) and IITP (MSIT 2019-0-01371). This work has been performed using resources provided by the Cambridge Tier-2 system operated by the University of Cambridge Research Computing Service (www.hpc.cam.ac.uk) funded by EPSRC Tier-2 capital grant (EP/T022159/1). HPC access was additionally funded by an EPSRC research infrastructure grant to F.M.. We are grateful to all the study participants and the Wolfson Brain Imaging Centre for their support. For the purpose of open access, the author has applied a Creative Commons Attribution (CC BY) licence to any Author Accepted Manuscript version arising from this submission.

## 7 AUTHOR CONTRIBUTIONS

BS, FM, ML and NS designed the study. AG and IS collected the data. AG, JO, FM, PM analysed the data. BS, FM and PM wrote the article.

