## supplementary information and analyses for "Enhanced behavioural and neural sensitivity to punishments in chronic pain and fatigue"

### SUPPLEMENTARY METHODS AND RESULTS

#### CLINICAL AND DEMOGRAPHIC DATA

Clinical and demographic data of the 57 participants who completed the full study are reported in Table S1. As expected, patients scored higher than controls on the Brief Pain Inventory (severity score: t55=11.188, p<0.001, Cohen's d=1.860, BF10=$4.495x10^{(}+12)$; interference score: t55=8.074, p<0.001, BF10=$9.935x10^{(}+7)$), McGill Pain Questionnaire (t55=7.807, p<0.001, Cohen's d=2.068, BF10=$3.873x10^{(}+7)$). Patients reported more fatigue (FACIT scale: t55=-9.055, p<0.001, Cohen's d=-2.399, BF10=3.132e+9) and poorer sleep quality than controls (PROMIS sleep scale: t55=4.784, p<0.001, Cohen's d=1.268, BF10=1302.373). This is paralleled by higher mean ratings of pain (t55=12.299, p<0.001, Cohen's d=3.259, BF10=1.650e+14) and fatigue (t55=8.672, p<0.001, Cohen's d=2.298, BF10=8.177e+8), and lower ratings of sleep quality (t55=2.948, p=0.005, Cohen's d=0.788, BF10=8.667), collected using diaries over a continuous period of 5 days, relative to controls. Average levels of pain and fatigue were highly correlated across participants (Pearson's r=0.84, BF10=$2.210x10^{1}3$). Interestingly, patients adopted higher severity ratings to describe their fatigue levels relative to their pain levels (mean fatigue diary rating ± SD, 5.461 ± 1.765; mean pain diary rating ± SD, 4.804 ± 1.698; t28=2.780, p=0.010, Cohen's d=0.516, BF10=4.710). Unsurprisingly, patients consumed more caffeine per day (t55=3.444, p=0.001, Cohen's d=0.913, BF10=28.578), but comparable units of alcohol than controls (t55=0.221, p=0.826, Cohen's d=0.059, BF10=0.274). Finally, patients reported greater depression symptoms at the Beck Depression Inventory (mean score ± SD, 8.429±5.295; group difference, t55=6.961, p<0.001, Cohen's d=1.860, BF10=1.798e+6) and generalised anxiety symptoms at the GAD7 test (mean score ± SD, 7.857±4.897; group difference, t55=6.560, p<0.001, Cohen's d=1.753, BF10=448796.707). Overall, we found strong evidence for associations between pain and fatigue questionnaires (BPI, McGIll and FACIT) and self-reported measures of depression and generalised anxiety (all correlations $p < 0.001$).

#### Table S1. Demographic and clinical data of the participants who completed the study.

| ID | Group | Gender | Age | Diagnosis | DAS/DAPSA score | BPI severity | BPI interference | McGIll | Diary avg pain | Facit | Diary avg fatigue | BDI score | GAD score | Promis (raw score) | Diary avg sleep | CRP | ESR | RheumatoidFactor | IFNγ | IL-10 | IL-12p70 | IL-13 | IL-1β | IL-2 | IL-4 | IL-6 | IL-8 | TNFα |
|---|---|---|---|---|---|---|---|---|---|---|---|---|---|---|---|---|---|---|---|---|---|---|---|---|---|---|---|---|
| P08 | Control | Male | 60 | n/a | 0.56 | 0.25 | 0.00 | 0 | 0.43 | 49 | 2.05 | 0 | 0 | 14 | 3.73 | <4 | 2 | <10 | 3.33 | 0.21 | 0.33 | 0.91 | 0.13 | 0.46 | <0.1 | 0.81 | 4.87 | 1.65 |
| P10 | Control | Female | 66 | n/a | 2.26 | 0 | 0.00 | 0 | 0.45 | 52 | 1.91 | 0 | 0 | 10 | 0.67 | <4 | 16 | <10 | 1.88 | 0.37 | 0.87 | 1.94 | <0.1 | 0.80 | <0.1 | 0.64 | 3.04 | 0.66 |
| P12 | Control | Female | 43 | n/a | 1.46 | 2.75 | 0.29 | 8 | 1.33 | 28 | 3.93 | 3 | 1 | 25 | 7.29 | <4 | 6 | 10 | 4.23 | 0.22 | <0.2 | <0.8 | 0.10 | 0.52 | <0.1 | 0.32 | 9.12 | 2.46 |
| P13 | Control | Male | 55 | n/a | 1.97 | 0.25 | 0.43 | 0 | 0.64 | 46 | 0.64 | 1 | 0 | 11 | 1.29 | <4 | 11 | <10 | 18.08 | 0.20 | <0.2 | <0.8 | <0.1 | 0.65 | <0.1 | 3.24 | 3.76 | 1.86 |
| P14 | Control | Female | 65 | n/a | 2.02 | 0 | 0.00 | 0 | 0.00 | 43 | 0.67 | 1 | 4 | 15 | 1.43 | <4 | 18 | <10 | 2.96 | 0.08 | 0.41 | <0.8 | <0.1 | 1.70 | <0.1 | 0.45 | 6.19 | 2.24 |
| P15 | Control | Female | 62 | n/a | 1.27 | 0 | 0.00 | 0 | 0.00 | 47 | 3.54 | 0 | 4 | 23 | 6.71 | <4 | 6 | <10 | 13.94 | 0.57 | 0.56 | 1.15 | 0.38 | 0.53 | <0.1 | 1.75 | 9.23 | 4.37 |
| P20 | Control | Female | 50 | n/a | 0.49 | 0 | 0.00 | 0 | 2.26 | 47 | 2.36 | 0 | 1 | 28 | 5.14 | <4 | 2 | 10 | 7.70 | 0.09 | <0.2 | <0.8 | <0.1 | <0.2 | <0.1 | 0.51 | 6.27 | 1.38 |
| P23 | Control | Male | 51 | n/a | 1.53 | 0.75 | 0.29 | 4 | 0.71 | 43 | 1.14 | 0 | 2 | 21 | 4.86 | <4 | 4 | 11 | 6.29 | 0.32 | 0.47 | 1.03 | 0.16 | 0.43 | <0.1 | 0.79 | 4.36 | 31.99 |
| P32 | Control | Male | 51 | n/a | 1.36 | 0 | 0.00 | 0 | 0.00 | 52 | 2.07 | 0 | 0 | 12 | 1.43 | <4 | 7 | <10 | 3.04 | 0.25 | 0.53 | <0.8 | <0.1 | 2.57 | <0.1 | 0.55 | 3.61 | 1.75 |
| P37 | Control | Female | 50 | n/a | 1.46 | 0 | 0.00 | 0 | 0.00 | 52 | 1.23 | 0 | 0 | 16 | 3.29 | <4 | 8 | <10 | 4.15 | 0.20 | <0.2 | 0.85 | <0.1 | 0.41 | <0.1 | 0.96 | 5.77 | 2.84 |
| P43 | Control | Female | 33 | n/a | 1.25 | 1 | 0.29 | 0 | 1.20 | 49 | 1.73 | 5 | 2 | 23 | 3.13 | <4 | 4 | 11 | 7.18 | 0.52 | 0.47 | 1.28 | 0.50 | 0.63 | <0.1 | 1.11 | 8.48 | 2.87 |
| P46 | Control | Male | 42 | n/a | 1.39 | 0 | 0.00 | 0 | 0.00 | 50 | 2.13 | 1 | 0 | 23 | 3.50 | <4 | 6 | <10 | 2.56 | 0.25 | 0.20 | <0.8 | 0.11 | 0.24 | <0.1 | 0.63 | 3.45 | 1.23 |
| P47 | Control | Female | 67 | n/a | 1.2 | 1 | 0.43 | 4 | 0.64 | 46 | 1.64 | 4 | 3 | 25 | 3.33 | <4 | 5 | 11 | n/a | n/a | n/a | n/a | n/a | n/a | n/a | n/a | n/a | n/a |
| P49 | Control | Female | 38 | n/a | 1.54 | 0 | 0.00 | 0 | 0.17 | 52 | 1.92 | 0 | 0 | 14 | 7.29 | <4 | 9 | <10 | 14.92 | 0.55 | 0.48 | 1.68 | 0.47 | 0.32 | <0.1 | 2.43 | 8.16 | 23.14 |
| P50 | Control | Male | 69 | n/a | n/a | 0 | 0.00 | 0 | 0.00 | 45 | 3.21 | 0 | 0 | 14 | 1.00 | n/a | n/a | n/a | 2.38 | 0.29 | <0.2 | 0.98 | <0.1 | <0.2 | <0.1 | 0.68 | 4.95 | 2.03 |
| P55 | Control | Female | 68 | n/a | 2.33 | 1 | 0.00 | 2 | 1.53 | 48 | 2.80 | 5 | 5 | 16 | 3.29 | <4 | n/a | <10 | 4.04 | 0.23 | 0.40 | 0.81 | 0.15 | 0.44 | <0.1 | 1.22 | 4.52 | 2.50 |
| P58 | Control | Female | 59 | n/a | n/a | 0 | 0.00 | 0 | 0.60 | 52 | 0.73 | 4 | 6 | 14 | 1.00 | n/a | n/a | n/a | 2.65 | 0.23 | 0.61 | 1.31 | <0.1 | 0.34 | <0.1 | 0.86 | 2.49 | 1.90 |
| P59 | Control | Male | 65 | n/a | 1 | 0 | 0.00 | 0 | 0.18 | 50 | 1.45 | 0 | 0 | 16 | 2.67 | <4 | 4 | <10 | 10.30 | 0.18 | 0.33 | <0.8 | <0.1 | 0.33 | <0.1 | 1.28 | 3.31 | 2.65 |
| P60 | Control | Female | 64 | n/a | 1.13 | 0 | 0.00 | 0 | 0.07 | 49 | 0.29 | 1 | 1 | 16 | 4.29 | <4 | 5 | <10 | 12.56 | 0.37 | 0.46 | 1.97 | 0.16 | 0.65 | <0.1 | 1.65 | 5.06 | 2.05 |
| P68 | Control | Female | 58 | n/a | 0.96 | 0 | 0.00 | 0 | 0.00 | 42 | 1.25 | 0 | 0 | 16 | 6.50 | <4 | n/a | <10 | 2.07 | 0.41 | 0.28 | <0.8 | <0.1 | 0.34 | <0.1 | 0.55 | 3.22 | 1.96 |
| P71 | Control | Female | 42 | n/a | 1.46 | 0 | 0.00 | 0 | 0.00 | 50 | 0.80 | 2 | 2 | 17 | 2.00 | <4 | 8 | 10 | 9.96 | 0.12 | 0.47 | 1.37 | <0.1 | 1.56 | <0.1 | 0.55 | 6.82 | 2.05 |
| P72 | Control | Male | 54 | n/a | 0.56 | 0 | 0.00 | 0 | 0.00 | 48 | 4.29 | 0 | 0 | 20 | 5.43 | <4 | 2 | <10 | 5.68 | 0.74 | 0.79 | <0.8 | 0.18 | 0.45 | <0.1 | 0.48 | 6.67 | 1.91 |
| P74 | Control | Male | 44 | n/a | 1.27 | 0 | 0.00 | 0 | 0.00 | 51 | 1.57 | 0 | 0 | 12 | 2.71 | <4 | 5 | <10 | 2.77 | 0.24 | 0.32 | 2.30 | <0.1 | 0.34 | <0.1 | 1.36 | 4.00 | 1.79 |
| P75 | Control | Female | 61 | n/a | 2.02 | 1.5 | 0.57 | 9 | 2.17 | 48 | 0.83 | 2 | 0 | 14 | 1.14 | <4 | 12 | <10 | 8.20 | 0.32 | <0.2 | <0.8 | 0.10 | 0.40 | <0.1 | 1.32 | 4.93 | 1.71 |
| P76 | Control | Female | 46 | n/a | 1.13 | 0 | 0.00 | 0 | 0.00 | 45 | 5.47 | 2 | 2 | 28 | 5.43 | <4 | 5 | <10 | 7.61 | 0.32 | 0.46 | 0.96 | 0.18 | 0.42 | <0.1 | 2.77 | 4.11 | 2.02 |
| P78 | Control | Male | 39 | n/a | 0.97 | 0 | 0.00 | 0 | 2.10 | 50 | 2.70 | 1 | 2 | 21 | 4.00 | <4 | 4 | <10 | 2.95 | 0.71 | 0.58 | 1.14 | <0.1 | 0.55 | <0.1 | 0.45 | 4.47 | 39.77 |
| P79 | Control | Female | 48 | n/a | 1.36 | 0 | 0.00 | 0 | 0.00 | 47 | 1.00 | 0 | 4 | 23 | 4.86 | <4 | 7 | <10 | 16.39 | 0.28 | 0.22 | <0.8 | <0.1 | 0.46 | <0.1 | 0.67 | 9.94 | 3.81 |
| P80 | Control | Female | 57 | n/a | n/a | 0 | 0.00 | 0 | 0.00 | 52 | 0.00 | 0 | 0 | 22 | 2.43 | n/a | n/a | n/a | 4.78 | 0.33 | 2.23 | 1.06 | 0.16 | <0.2 | <0.1 | 0.73 | 5.26 | 18.17 |
| P01 | Patient | Female | 52 | RA | 4.95 | 6.5 | 4.29 | 23 | 7.31 | 27 | 7.31 | 2 | 4 | 28 | 5.71 | <4 | 28 | n/a | 31.74 | 2.67 | 3.62 | 6.36 | 1.67 | 0.85 | 0.30 | 8.65 | 12.59 | 4.51 |
| P02 | Patient | Female | 56 | RA | 4.96 | 5.25 | 6.57 | 13 | 5.08 | 36 | 6.27 | 7 | 16 | 33 | 5.88 | <4 | 5 | 36 | 3.17 | 0.27 | 0.74 | <0.8 | <0.1 | 0.56 | <0.1 | 1.76 | 3.92 | 1.97 |
| P03 | Patient | Female | 50 | RA | 3.5 | 3.25 | 2.29 | 9 | 4.00 | 38 | 3.73 | 6 | 8 | 29 | 2.00 | <4 | 11 | <10 | 2.25 | 0.29 | 0.36 | 0.95 | <0.1 | 0.29 | <0.1 | 1.10 | 5.68 | 1.69 |
| P05 | Patient | Male | 60 | RA | 2.62 | 2.25 | 2.43 | 4 | 1.99 | 29.55 | 4.55 | 4 | 5 | 20 | 3.70 | <4 | 16 | 56 | 10.36 | 1.36 | 0.57 | 1.31 | <0.1 | 0.39 | <0.1 | 12.08 | 7.03 | 3.52 |
| P07 | Patient | Female | 36 | RA | 6.94 | 7 | 7.00 | 20 | 7.10 | 12 | 7.00 | 19 | 15 | 33 | 5.50 | 34 | 19 | 101 | 4.46 | 0.28 | 0.53 | <0.8 | 0.30 | 0.46 | <0.1 | 0.54 | 5.17 | 2.62 |
| P16 | Patient | Female | 63 | RA | 2.66 | 1.25 | 1.14 | 2 | 2.00 | 33 | 2.93 |  |  | 19 | 2.43 | <4 | 5 | <10 | 3.20 | 0.53 | 0.29 | <0.8 | <0.1 | 0.37 | <0.1 | 6.50 | 5.31 | 2.46 |
| P17 | Patient | Female | 55 | RA | 1.63 | 1 | 0.00 | 2 | 2.33 | 49 | 2.93 | 2 | 6 | 17 | 3.14 | <4 | 9 | 12 | 3.20 | 0.12 | 0.38 | <0.8 | <0.1 | 0.31 | <0.1 | 0.67 | 5.54 | 1.90 |
| P18 | Patient | Female | 37 | RA | 3.75 | 4.75 | 2.43 | 22 | 4.87 | 28 | 2.07 | 5 | 9 | 22 | n/a | <4 | 9 | 25 | 3.36 | 0.66 | 0.25 | <0.8 | <0.1 | 0.33 | <0.1 | 0.37 | 5.51 | 2.05 |
| P19 | Patient | Female | 59 | RA | 3.28 | 3 | 2.00 | 13 | 3.00 | 40 | 2.53 | 1 | 2 | 17 | 7.86 | n/a | n/a | n/a | 2.62 | 0.14 | <0.2 | <0.8 | 0.26 | <0.2 | <0.1 | 0.25 | 2.71 | 1.40 |
| P24 | Patient | Female | 54 | RA | 3.27 | 2.5 | 1.57 | 6 | 3.53 | 41 | 3.33 | 10 | 5 | 19.43 | 5.43 | 30 | 12 | 80 | 6.66 | 0.27 | <0.2 | 1.04 | 0.12 | 0.43 | <0.1 | 3.07 | 3.18 | 2.01 |
| P27 | Patient | Female | 56 | RA | 4.74 | 1.5 | 0.43 | 4 | 2.40 | 23 | 4.33 | 5 | 2 | 26 | 6.43 | <4 | 11 | 10 | 9.46 | 0.21 | 0.96 | 1.48 | 0.26 | 0.58 | <0.1 | 0.77 | 4.23 | 2.65 |
| P28 | Patient | Female | 65 | RA | 4.19 | 4.75 | 5.57 | 11 | 4.73 | 29.25 | 6.80 | 12 | 18 | 25 | 4.63 | <4 | 8 | 44 | 5.20 | 0.28 | 0.77 | <0.8 | <0.1 | 0.62 | <0.1 | 0.52 | 3.20 | 1.90 |
| P29 | Patient | Female | 37 | RA | 4.4 | 2.5 | 1.29 | 12 | 3.42 | 33 | 4.68 | 5 | 6 | 25 | 6.29 | <4 | 13 | 67 | 5.71 | 0.31 | 0.58 | 0.81 | <0.1 | 0.60 | <0.1 | 12.19 | 13.82 | 3.02 |
| P33 | Patient | Female | 47 | RA | 4.26 | 3.5 | 1.57 | 3 | 4.60 | 36 | 5.80 | 6 | 6 | 14 | 3.86 | <4 | 11 | 17 | 2.17 | 0.26 | 0.21 | 3.89 | <0.1 | <0.2 | <0.1 | 0.34 | 5.30 | 2.18 |
| P34 | Patient | Female | 57 | RA | 6.75 | 6 | 4.29 | 17 | 7.80 | 7 | 7.92 | 6 | 15 | 28 | 5.67 | 8 | 23 | 252 | 2.73 | 0.13 | 0.22 | <0.8 | <0.1 | 0.39 | <0.1 | 2.80 | 3.51 | 1.74 |
| P35 | Patient | Female | 54 | RA | 6 | 4.25 | 8.43 | 21 | 4.47 | 16 | 4.07 | 22 | 14 | 17 | 1.60 | <4 | 18 | 19 | 10.89 | 0.15 | 0.26 | <0.8 | <0.1 | 0.21 | <0.1 | 8.75 | 9.06 | 2.64 |
| P38 | Patient | Female | 60 | RA | 5.91 | 5 | 6.57 | 17 | 6.09 | 22 | 5.67 | 9 | 5 | 22 | 5.20 | <4 | 8 | 143 | 6.11 | 0.29 | 0.39 | 1.09 | <0.1 | 0.38 | <0.1 | 1.30 | 3.28 | 1.99 |
| P40 | Patient | Female | 57 | RA | 7.18 | 5.75 | 5.71 | 4 | 5.69 | 34 | 7.00 | 5 | 9 | 38 | 7.17 | 16 | 41 | 73 | 4.61 | 0.29 | 0.77 | <0.8 | 0.14 | <0.2 | <0.1 | 0.57 | 4.39 | 2.77 |
| P42 | Patient | Female | 53 | PsA | 5.63 | 3.75 | 4.57 | 10 | 3.85 | 33 | 4.23 | 14 | 6 | 23 | 2.33 | <4 | 7 | 11 | 10.22 | 0.73 | <0.2 | <0.8 | <0.1 | 0.37 | <0.1 | 1.61 | 6.44 | 2.16 |
| P44 | Patient | Female | 35 | RA | 3.51 | 6 | 7.00 | 17 | 3.36 | 31 | 4.77 | 11 | 5 | 23 | 4.57 | 17 | 12 | <10 | 4.54 | 0.22 | 0.31 | <0.8 | <0.1 | 0.27 | <0.1 | 0.50 | 4.14 | 1.63 |
| P54 | Patient | Female | 52 | RA | 5.46 | 6.75 | 8.43 | 20 | 4.77 | 21 | 7.77 | 10 | 16 | 38 | 7.00 | 21 | 18 | <10 | 4.08 | 0.64 | <0.2 | <0.8 | 0.11 | 0.29 | <0.1 | 0.86 | 3.46 | 2.35 |
| P56 | Patient | Female | 45 | RA | 5.66 | 6.75 | 10.00 | 26 | 6.47 | 14 | 5.73 | 13 | 13 | 32 | 5.00 | 22 | 35 | 368 | 2.94 | 0.33 | 2.02 | <0.8 | <0.1 | 0.99 | <0.1 | 0.48 | 3.62 | 1.88 |
| P57 | Patient | Female | 54 | RA | 4.76 | 3.25 | 5.00 | 15 | 4.94 | 27 | 6.38 | 4 | 0 | 24 | 4.50 | 19 | 21 | <10 | 3.67 | 0.10 | <0.2 | <0.8 | <0.1 | <0.2 | <0.1 | 0.95 | 3.95 | 2.00 |
| P61 | Patient | Male | 45 | PsA | 3.2 | 5.25 | 4.43 | 21 | 5.60 | 32 | 8.30 | 8 | 10 | 39 | 9.20 | <4 | 5 | <10 | 2.28 | 0.97 | <0.2 | <0.8 | 0.22 | 0.86 | <0.1 | 2.23 | 6.00 | 2.65 |
| P64 | Patient | Female | 49 | RA | 3.28 | 4.25 | 2.29 | 4 | 5.46 | 32 | 6.31 | 4 | 3 | 24 | 6.29 | n/a | n/a | n/a | <1.7 | 0.70 | 0.83 | <0.8 | <0.1 | 0.22 | <0.1 | 0.97 | 2.68 | 1.35 |
| P65 | Patient | Female | 62 | PsA | 5 | 6.25 | 6.43 | 11 | 7.64 | 25 | 6.21 | 6 | 5 | 31 | 6.14 | 29 | 47 | <10 | 4.40 | 0.24 | <0.2 | <0.8 | 0.11 | 0.45 | <0.1 | 1.57 | 5.66 | 2.61 |
| P66 | Patient | Female | 62 | RA | 4.24 | 3.75 | 6.43 | 24 | 5.00 | 3 | 6.60 | 16 | 5 | 23 | 3.50 | <4 | 9 | 114 | 2.25 | 0.27 | 0.62 | 0.93 | 0.23 | 0.93 | <0.1 | 0.45 | 4.03 | 1.65 |
| P69 | Patient | Female | 39 | RA | 6.93 | 7 | 8.43 | 19 | 7.50 | 11 | 8.00 | 16 | 6 | 36 | 6.67 | <4 | 32 | <10 | 4.97 | 0.22 | 0.77 | 1.52 | <0.1 | 0.56 | <0.1 | 4.87 | 4.15 | 2.16 |
| P73 | Patient | Female | 45 | PsA | 4.12 | 3 | 0.71 | 2 | 4.33 | 34 | 5.13 | 8 | 6 | 26 | 4.29 | 18 | 25 | 10 | 6.72 | 0.38 | 0.35 | <0.8 | 0.14 | 0.59 | <0.1 | 1.15 | 4.76 | 27.27 |

### ADDITIONAL TASK RESULTS

Patients won a comparable amount of money (net £4.62±2.50) than controls (net £5.18±2.53) in the 4-armed bandit task (t55=0.837, p=0.406, Cohen's d=0.222, BF10=0.359). Reaction times were comparable between groups (t55=0.794, p=0.430, Cohen's d=0.210, BF10=0.349).

Patients and controls were similarly likely to switch choice, irrespective of the outcome (t55=0.528, p=0.6, Cohen's d=0.140, BF10=0.301). However, patients were overall slower at making a choice than controls (p=0.013) and we found a nominally-significant interaction between group and choice type (2 levels: repeated, switched; p=0.027); the choice to switch was slower in patients than controls (t55=2.478, p=0.016, Cohen's d=0.657, BF10=3.243), whereas the choice to stick to the previous decision had similar latency in the two groups (t55=1.484, p=0.144, Cohen's d=0.393, BF10=0.667).

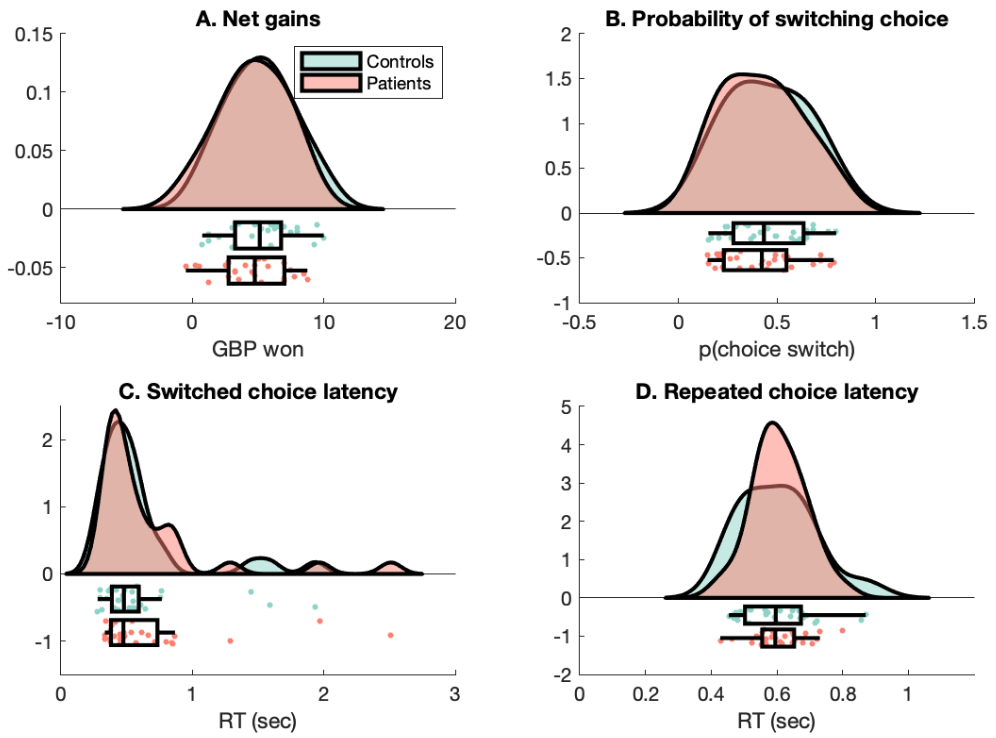


**Figure S1. Task performance.** A. Net gains (GBP); B. Probability of switching choice irrespective of the outcome; C. Latency of the switched choice; D. Latency of the repeated choice in controls (green) and patients (red). For each variable, the probability density functions, scatterplot of individual data and boxplot are shown.

We evaluated whether the frequency of choice switches was affected by the outcome of the preceding trial (4 levels: reward only, punishment only, simultaneous reward and punishment, no outcome) and by the diagnosis (2 levels: inflammatory arthritis, healthy controls). A mixed-effects ANOVA provided evidence for an effect of condition (F1.45,79.75=110.824, p<0.001, eta2=0.637), but no evidence for main effect of group (F1,55=0.907, p=0.345, eta2=0.016) nor interaction (F1.45,79.75=0.517, p=0.540, eta2=0.003). Irrespective of their group, participants were more likely to change their choice following a trial that yielded either a loss or no outcome than in trials that yielded either a win or a simultaneous win and loss (all ps < 0.001, Cohen's d > 1.5). Moreover, the frequency of choice switches was similar after a loss and no outcome (p=0.062, Cohen's d=0.253); participants switched choices less frequently after affording a win rather than an ambiguous outcome (p=0.017, Cohen's d=0.360). Finally, the total frequency of choice switches were comparable across groups (t55=0.952, p=0.345, Cohen's d=0.252, BF10=0.391).

### SLEEP ACTIGRAPHY

We used ActiLife v6.13.3 to extract sleep measures. Automatic detection of sleep periods was implemented with Tudor-Locke automatic sleep period detection. If there was a discrepancy of >2 hours between the automatic detected timing and the self-documented bedtime or out-of-bed time, the self-documented times would be entered manually as sleep periods. The Cole-Kripke algorithm was applied to the data to determine sleep parameters (Cole et al., 1992). Data from 5 participants (3 patients) with <5 nights of recording were excluded from further analyses, leaving 52 participants for analysis (25 patients).

Controls and patients showed comparable results in the sleep parameters, total sleep time (t(50)=1.597, p=.117, Cohen's d=0.443, BF10=0.788), movement index (t(50=-1.684, p=0.098, Cohen's d=0.467, BF10=0.884), sleep fragmentation (t(50)=-1.648, p=0.106, Cohen's d=0.457, BF10=0.842), sleep latency (t(50)=-0.296, p=0.768, Cohen's d=0.082, BF10=0.289), sleep efficiency (t(50)=0.794, p=0.431, Cohen's d=0.220, BF10=0.361) and wake-after-sleep-onset (t(50)=-0.171, p = 0.865, Cohen's d=0.047, BF10=0.282).

### DAILY ACTIVITY

We used ActiLife v6.13.3 to extract daily activity measures, implementing the default wear time validation and (Freedson et al., 1998) algorithm to detect activity bouts. Days with < 1080 epochs (i.e., 18 hours) of data were excluded from the analyses (154 days removed in total). Similar to the sleep analysis, any participants with < 5 days of recording were removed from further analysis (N=5, 2 patients), leaving 52 participants for analysis (26 patients).

Patients and controls showed comparable results in the activity parameters, with mean daily steps (mean ±SD, patients: 12114.340 ±4323.304, controls: 14191.425 ± 3714.191; t(50)= 1.858, p=.069, Cohen's d=0.515, BF10=1.131), proportion spent in sedentary activity (mean ± SD, patients: 0.536 ± 0.108, controls: 0.498 ±0.098; t(50)=-1.347, p=.184, Cohen's d=0.374, BF10=0.585), proportion spent in light activity (mean ± SD, patients: 0.355 ± 0.083, controls: 0.371 ± 0.072; t(50)=0.768, p=.446, Cohen's d=0.213, BF10=0.355), proportion spent in moderate activity (mean ± SD, patients: 0.109 ± 0.062, controls: 0.131 ± 0.043; t(50)=1.487, p=.143, Cohen's d=0.413, BF10=0.688), neither group spending any time in vigorous activity, number of bouts (mean ± SD, patients: 8.371 ± 5.703, controls: 9.667 ± 4.585; t(50)=0.903, p=.371, Cohen's d=0.251, BF10=0.389), total time of bouts (mean ± SD, patients: 137.213 ± 100.688, controls: 162.494 ±77.400; t(50)=1.015, p=.315, Cohen's d=0.282, BF10=0.425), and total activity bout count (mean ± SD, patients: 462028.583 ± 350630.645, controls: 596035.778 ± 253689.269; t(50=1.579, p=.121, Cohen's d=0.438, BF10=0.770).

### SUPPLEMENTARY MODELLING ANALYSES

#### 8.1 Parameter recovery

To asses the reliability of our modelling analysis (Wilson and Collins, 2019), for each model we performed parameter recovery analysis, where we simulated participants' responses using newly drawn individual-level parameters from the group-level distributions, for each group and for a pooled dataset.

We then fit the same model to the simulated data and calculated Pearson correlation coefficients $r$ between the generated and estimated individual-level parameters. We recovered each individual parameter (for each member of the population dataset) 50 times and calculated the mean and SD of the correlation between the true and recovered parameter values. See results in S2-S4. The higher the coefficient $r$, the more reliable the estimates are, which can be categorised as: poor (if r <0.5); fair (if 0.5< r <0.75); good (0.75< r <0.9); excellent (if r >0.9) (White et al., 2018).

Moreover, to assess the number of simulations needed, we calculated the average error of the parameter recovery correlation (and its error) for each model, parameter, and dataset as a function of increasing number of simulation, as plotted in Figure S2. The average was obtained from the 1000 randomly chosen permutations of different simulations at each $n$ (out of 50).

**Table S2.** Pearson correlation coefficients (SD) for parameter recovery analysis for each parameter from five RL models for a pooled dataset.

| Model ID | LR1 | LR2 | R | P | $\xi$ | d |
|---|---|---|---|---|---|---|
| 1 | 0.91 (0.03) | 0.75 (0.09) | 0.87 (0.04) | 0.78 (0.06) | 0.67 (0.08) | 0.92 (0.02) |
| 2 | 0.90 (0.11) | 0.75 (0.16) | 0.83 (0.09) | 0.79 (0.13) | 0.61 (0.17) | |
| 3 | 0.92 (0.02) | 0.26 (0.11) | 0.85 (0.04) | 0.91 (0.03) | 0.65 (0.10) | 0.92 (0.02) |
| 4 | 0.95 (0.02) | 0.88 (0.04) | 0.88 (0.03) | 0.90 (0.03) | 0.70 (0.11) | |
| 5 | 0.95 (0.02) | 0.88 (0.03) | 0.94 (0.02) | 0.92 (0.03) | | |

**Table S3.** Pearson correlation coefficients (SD) for parameter recovery analysis for each parameter from five RL models for patients.

| Model ID | LR1 | LR2 | R | P | $\xi$ | d |
|---|---|---|---|---|---|---|
| 1 | 0.93 (0.03) | 0.78 (0.11) | 0.84 (0.05) | 0.61 (0.11) | 0.76 (0.09) | 0.93 (0.03) |
| 2 | 0.90 (0.11) | 0.73 (0.20) | 0.76 (0.16) | 0.73 (0.17) | 0.70 (0.19) | |
| 3 | 0.90 (0.06) | 0.20 (0.16) | 0.83 (0.06) | 0.90 (0.04) | 0.69 (0.10) | 0.91 (0.05) |
| 4 | 0.96 (0.02) | 0.91 (0.05) | 0.80 (0.07) | 0.81 (0.07) | 0.73 (0.11) | |
| 5 | 0.95 (0.03) | 0.91 (0.04) | 0.94 (0.03) | 0.86 (0.05) | | |

**Table S4.** Pearson correlation coefficients (SD) for parameter recovery analysis for each parameter from five RL models for healthy controls.

| Model ID | LR1 | LR2 | R | P | $\xi$ | d |
|---|---|---|---|---|---|---|
| 1 | 0.90 (0.05) | 0.14 (0.22) | 0.92 (0.04) | 0.92 (0.03) | 0.29 (0.16) | 0.90 (0.04) |
| 2 | 0.88 (0.11) | 0.72 (0.17) | 0.88 (0.11) | 0.88 (0.08) | 0.27 (0.25) | |
| 3 | 0.91 (0.04) | 0.22 (0.18) | 0.91 (0.03) | 0.93 (0.03) | 0.23 (0.18) | 0.89 (0.05) |
| 4 | 0.92 (0.03) | 0.83 (0.07) | 0.94 (0.04) | 0.94 (0.02) | 0.26 (0.17) | |
| 5 | 0.92 (0.03) | 0.80 (0.10) | 0.95 (0.02) | 0.94 (0.03) | | |

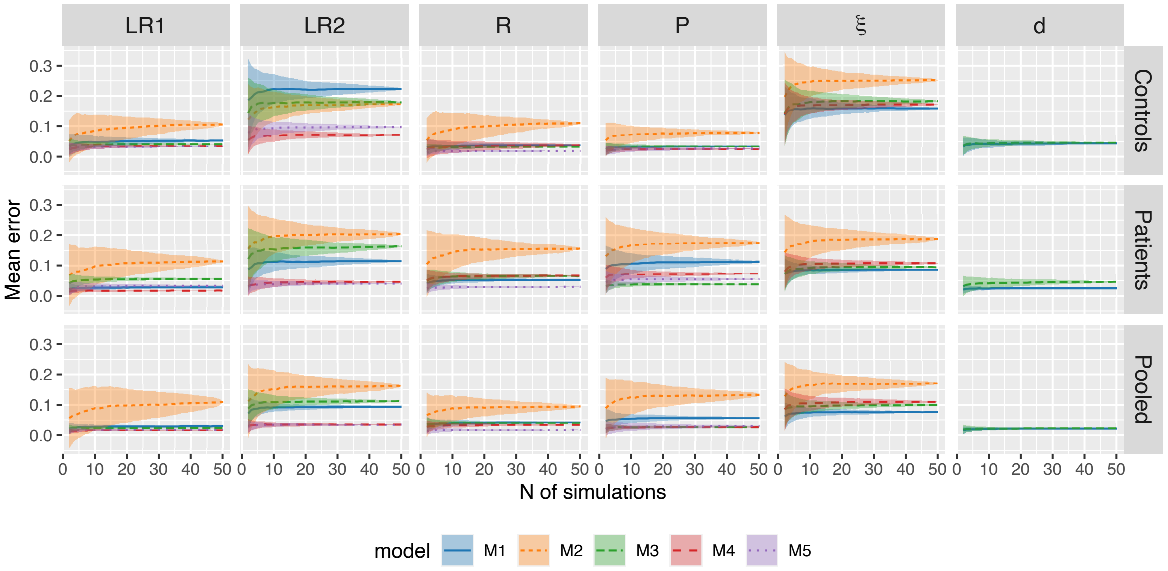


**Figure S2. Parameter recovery mean error** of the correlation for each model and each parameter. The average error is plotted as a function of simulation number averaged across 1000 permutations of 50 simulations.

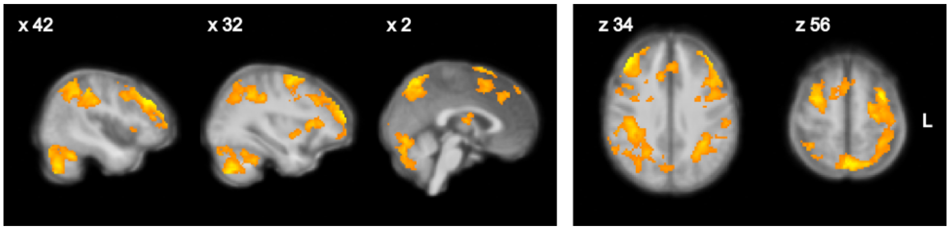


**Figure S3. Neural correlates of punishment prediction error shared across the arthritis and control groups.** Colorscale represents z scores ranging from 0 to 5, thresholded at z>2.6, p<0.05.

##### 8.2 Model recovery

We also performed model recovery analysis (Wilson and Collins, 2019), where we first simulated responses using each model and then fit each model to the model-specific dataset (50 simulations for each pair). We then counted the number of times a model fit the simulated data best (according to the LOOIC rule), effectively creating an $M \times M$ confusion matrix, where $M$ is the number of models. In the case where we have a diagonal matrix of ones, the models are perfectly recoverable and hence as reliable as possible. See the model recovery results for the pooled sample in Table S5.

**Table S5.** Confusion matrix from the model recovery analysis of five RL models for a pooled dataset.

| | | **fit with** | | | | |
|---|---|---|---|---|---|---|
| | | M1 | M2 | M3 | M4 | M5 |
| **generated** | M1 | 0.52 | 0.08 | 0.28 | 0.10 | 0.02 |
| | M2 | 0.22 | 0.08 | 0.22 | 0.34 | 0.14 |
| | M3 | 0.44 | 0.02 | 0.36 | 0.10 | 0.08 |
| | M4 | 0.22 | 0.14 | 0.22 | 0.22 | 0.20 |
| | M5 | 0.12 | 0.16 | 0.14 | 0.28 | 0.30 |

#### SUPPLEMENTARY IMAGING RESULTS

| Cluster ID | Voxels | P | Z peak | X | Y | Z |
|---|---|---|---|---|---|---|
| 2a | 16180 | 5.48e-30 | 5.62 | 42 | 36 | 34 |
| 2b | | | 5.34 | -32 | 0 | 56 |
| 2c | | | 4.97 | 2 | -64 | 54 |
| 2d | | | 4.91 | -44 | 20 | 38 |
| 2e | | | 4.83 | 4 | -72 | 50 |
| 2f | | | 4.83 | -4 | -66 | 56 |
| 1a | 8416 | 3.53e-19 | 4.81 | 32 | -70 | -44 |
| 1b | | | 4.76 | -34 | -48 | -30 |
| 1c | | | 4.59 | 42 | -66 | -38 |
| 1d | | | 4.58 | -40 | -56 | -44 |
| 1e | | | 4.57 | -26 | -66 | -40 |
| 1f | | | 4.57 | 20 | -78 | -20 |

**Table S6.** Activation clusters associated with punishment prediction error, common activity across patient and control groups. Coordinates in MNI space.

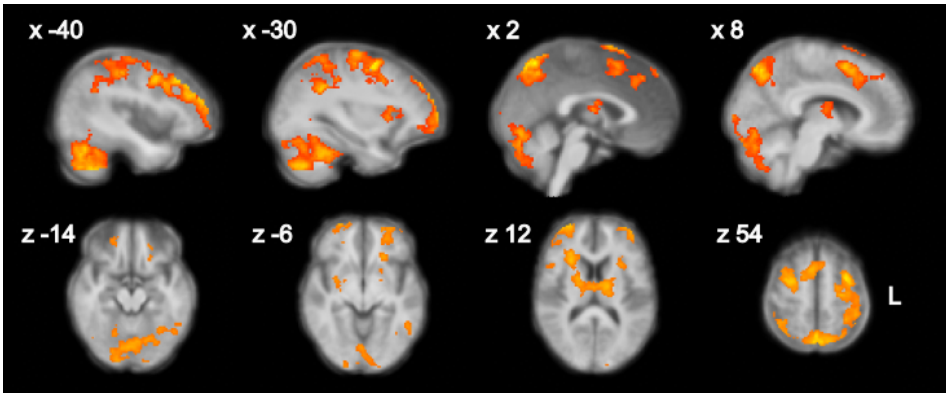


**Figure S4. Neural correlates of reward prediction error shared across the arthritis and control groups.** Colorscale represents z scores ranging from 0 to 5, thresholded at z>2.6, p<0.05.

| Cluster ID | Voxels | P | Z peak | X | Y | Z |
|---|---|---|---|---|---|---|
| 5a | 34881 | 0 | 7.08 | 12 | -86 | -14 |
| 5b | | | 6.59 | -26 | -76 | -16 |
| 5c | | | 6.52 | 26 | -82 | -16 |
| 5d | | | 6.38 | 22 | -84 | -16 |
| 5e | | | 6.35 | -26 | -84 | -18 |
| 5f | | | 6.22 | 26 | -92 | 2 |
| 4a | 2285 | 4.92E-17 | 4.47 | 2 | -12 | 12 |
| 4b | | | 3.83 | 8 | -12 | 12 |
| 4c | | | 3.79 | 8 | -22 | 10 |
| 4d | | | 3.79 | 28 | 4 | 24 |
| 4e | | | 3.78 | -10 | -12 | 12 |
| 4f | | | 3.72 | 28 | 10 | 18 |
| 3a | 1381 | 9.49E-12 | 4.38 | -30 | 2 | 54 |
| 3b | | | 3.9 | -38 | 0 | 24 |
| ] 3c | | | 3.71 | -40 | 4 | 24 |
| 3d | | | 3.7 | -42 | 4 | 34 |
| 3e | | | 3.7 | -38 | 28 | 24 |
| 3f | | | 3.53 | -38 | 30 | 38 |
| 2a | 712 | 5.36E-07 | 3.81 | -32 | 56 | -6 |
| 2b | | | 3.71 | -20 | 52 | -12 |
| 2c | | | 3.54 | -32 | 50 | -8 |
| 2d | | | 3.47 | -28 | 42 | -8 |
| 2e | | | 3.46 | -28 | 40 | -12 |
| 2f | | | 3.46 | -26 | 60 | 6 |
| 1a | 202 | 0.0346 | 3.98 | 10 | 56 | -12 |
| 1b | | | 3.94 | 12 | 58 | -8 |
| 1c | | | 3.6 | 8 | 60 | -6 |
| 1d | | | 3.35 | 26 | 60 | -6 |
| 1e | | | 3.09 | 26 | 56 | -8 |
| 1f | | | 2.92 | 20 | 52 | -10 |

**Table S7.** Activation clusters associated with reward prediction error, positive mean activity across patient and control groups. Coordinates in MNI space.

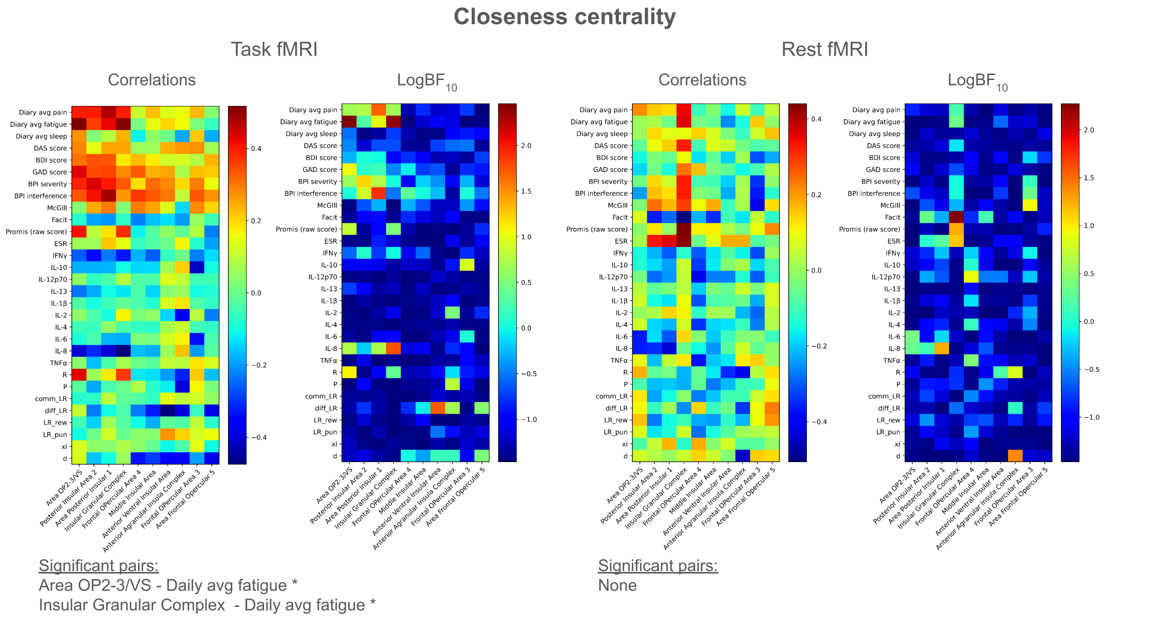


**Figure S5.** Closeness centrality correlation results. Please refer to the supplementary file with network analysis results for precise values of Pearson's R and LogBF10.

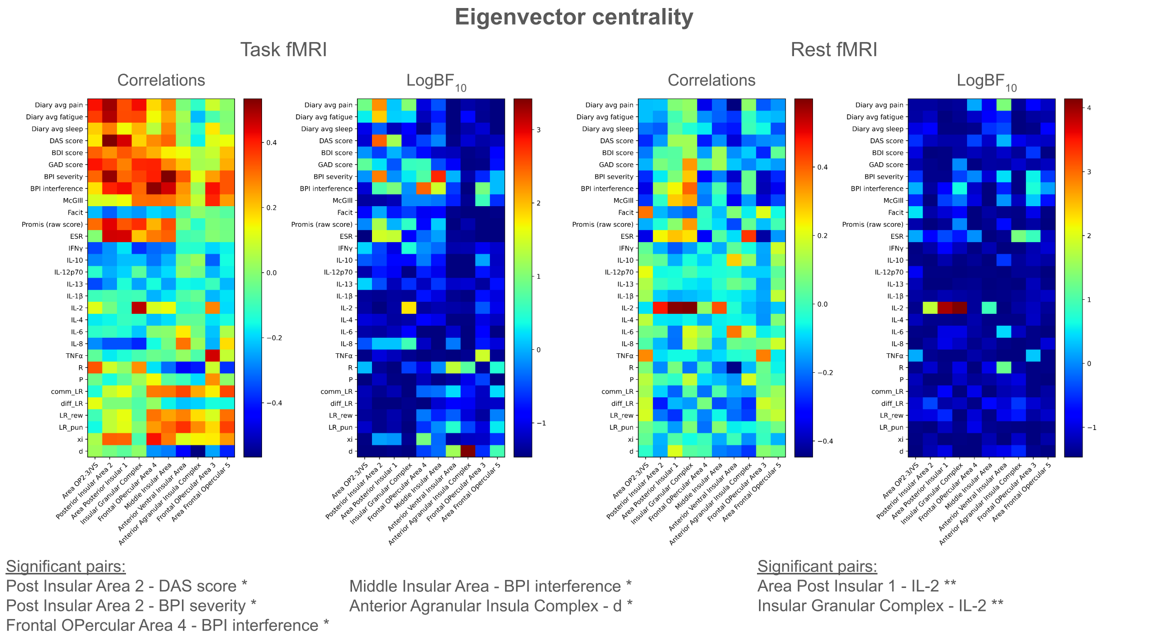


**Figure S6.** Eigenvector centrality correlation results. Please refer to the supplementary file with network analysis results for precise values of Pearson's R and LogBF10.

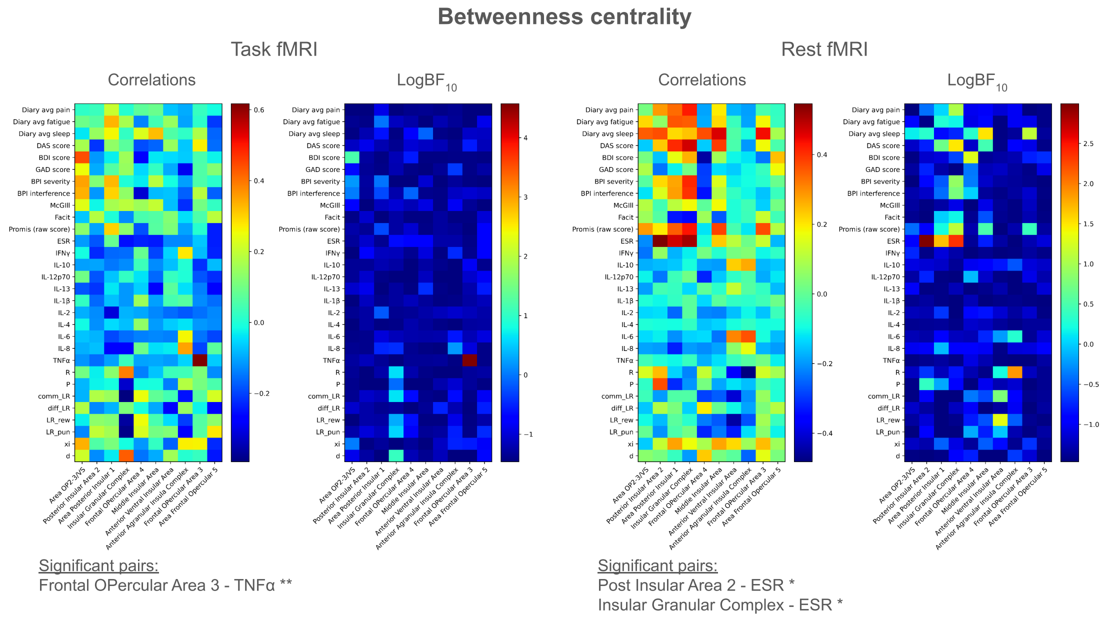


**Figure S7.** Betweenness centrality correlation results. Please refer to the supplementary file with network analysis results for precise values of Pearson's R and LogBF10.

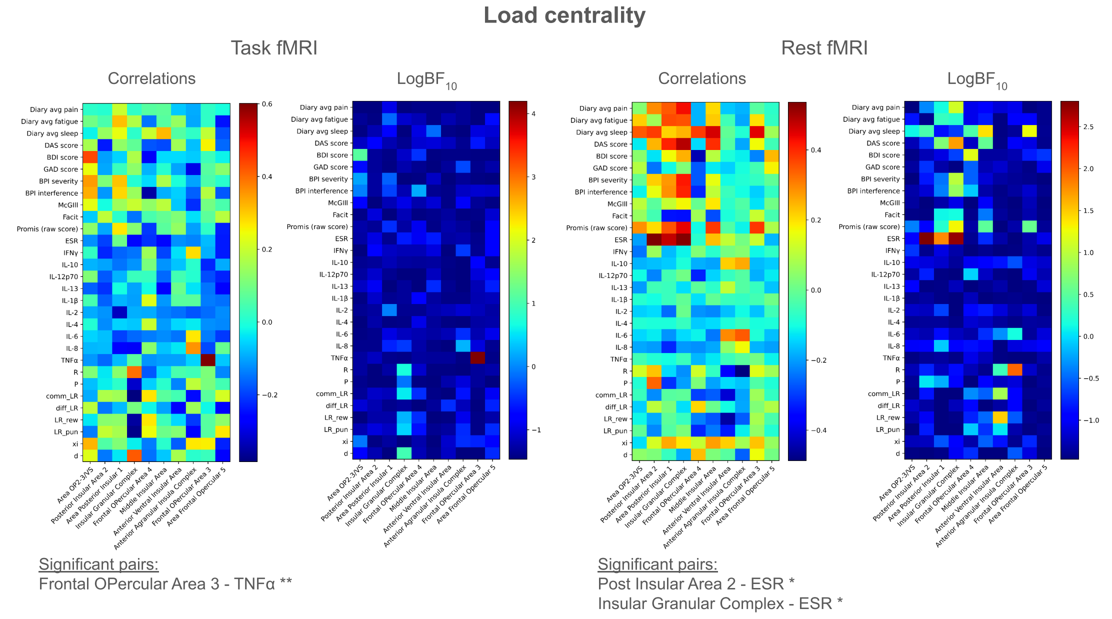


**Figure S8.** Load centrality correlation results. Please refer to the supplementary file with network analysis results for precise values of Pearson's R and LogBF10.
